# Probing pathophysiology of extracellular cGAMP with substrate-selective ENPP1

**DOI:** 10.1101/2021.05.04.442665

**Authors:** Jacqueline A. Carozza, Anthony F. Cordova, Yasmeen AlSaif, Volker Böhnert, Gemini Skariah, Lingyin Li

**Affiliations:** ChEM-H Institute, Stanford University, Stanford, CA 94301; Department of Chemistry, Stanford University, Stanford, CA 94301; Cancer Biology Program, Stanford University, Stanford, CA 94301; Department of Biology, Stanford University, Stanford, CA 94301; Department of Biochemistry, Stanford University, Stanford, CA 94301

## Abstract

The biology of the immune second messenger cGAMP depends on its cellular localization. cGAMP, which is synthesized in response to cytosolic double-stranded DNA, also exists in the extracellular space as a paracrine immunotransmitter that enhances the anticancer immune response. However, the role of extracellular cGAMP is unexplored outside of cancer due to a lack of tools to systemically manipulate it. The extracellular enzyme ENPP1, the only known hydrolase of cGAMP, is a promising target. However, because ENPP1 also degrades extracellular ATP, using genetic knockouts of ENPP1 to study extracellular cGAMP leads to confounding effects. Here we report the H362A point mutation in ENPP1, the dominant cGAMP hydrolase, which selectively abolishes ENPP1’s ability to degrade cGAMP, while retaining activity toward other substrates. H362 is not necessary for binding cGAMP or the catalytically-essential zinc atoms but instead supports the in-line reaction geometry. H362 is evolutionarily conserved down to bacteria, suggesting an ancient origin for extracellular cGAMP biology. *Enpp1^H362A^* mice do not display the systemic calcification seen in *Enpp1*^-/-^ mice, highlighting the substrate-specific phenotype of ENPP1. Remarkably, *Enpp1^H362A^* mice were resistant to HSV-1 infection, demonstrating the antiviral role of endogenous extracellular cGAMP. The ENPP1^H362A^ mutation is the first genetic tool to enable exploration of extracellular cGAMP biology in a wide range of tissues and diseases.

## Introduction

Cyclic-GMP-AMP (cGAMP) quickly became the intense focus of many biochemistry, immunology, and cell biology labs since its discovery eight years ago (Ablasser et al., 2013a; Diner et al., 2013; Wu et al., 2013). Its significance was immediately apparent, being a new addition to the relatively small group of known second messengers. cGAMP is synthesized by cells when they encounter double-stranded DNA (dsDNA) in the cytosol as a danger signal. At first, cGAMP was thought to be mainly produced during viral infections as an antiviral signal in innate immunity (Li et al., 2013b; Wu et al., 2013). Now, cGAMP signaling seems to have an impact in every disease the field has investigated: cGAMP signaling occurs in cancer cells (Bakhoum et al., 2018; Deng et al., 2014; Harding, 2017; Mackenzie et al., 2017; Woo et al., 2014), in neurodegenerative diseases (Sliter et al., 2018), during myocardial infarction (King et al., 2017), and in various autoimmune settings (Ahn et al., 2012; Gao et al., 2015). Furthermore, cGAMP-STING signaling has ancient evolutionary origins which have only just begun to be uncovered in bacteria and lower animals (Kranzusch et al., 2015; Morehouse et al., 2020; Whiteley et al., 2019). It has become clear that this innate immune second messenger has a broad and diverse range of physiological roles, just like the other second messengers discovered before it.

Just as our understanding of cGAMP’s disease implications has expanded over the years, so has our understanding of its cellular localization. cGAMP was originally thought to be an intracellular signal that is produced and sensed by STING within the same cell. Soon, research showed that cGAMP can spread from one cell to neighboring cells through gap junctions (Ablasser et al., 2013b; Chen et al., 2016; Schadt et al., 2019) or through secretion of exosomes that later fuse with other cells (Bridgeman et al., 2015; Gentili et al., 2015). In both of these cases cGAMP is sequestered from the extracellular compartment and is therefore an extension of intracellular cGAMP signaling. Like all second messengers, cGAMP signaling is subject to negative regulation by enzymatic degradation. However, unlike other second messengers, the cGAMP hydrolase ENPP1 is located outside the cell (Li et al., 2014), leading to our hypothesis that cGAMP can be secreted as an extracellular signal. We confirmed this new role for cGAMP as an extracellular immunotransmitter that is secreted by cancer cells and accumulates in the tumor microenvironment (Carozza et al., 2020a). It is taken up by surrounding cells via specific importers (Cordova et al., 2021; Lahey et al., 2020; Luteijn et al., 2019; Ritchie et al., 2019; Zhou et al., 2020), resulting in STING pathway activation. It is now clear that cancer-to-host extracellular cGAMP signaling activates anti-cancer innate immunity (Carozza et al., 2020a; Cordova et al., 2021; Li et al., 2020; Marcus et al., 2018; Schadt et al., 2019). We subsequently confirmed that ENPP1 only affects extracellular cGAMP concentrations and not intracellular cGAMP concentrations (Carozza et al., 2020a), highlighting the importance of cGAMP localization.

Does the importance of extracellular cGAMP extend beyond its role in cancer? What is the evolutionary conservation of extracellular cGAMP signaling? These are difficult questions to answer. Although it is possible to manipulate extracellular cGAMP in the confined tumor microenvironment (Carozza et al., 2020a; Cordova et al., 2021), it is technically challenging to systemically alter extracellular cGAMP. Since ENPP1 is the only known cGAMP hydrolase, it is an attractive target for manipulating extracellular cGAMP. However, it has been known for decades that ENPP1 also plays an essential role by degrading extracellular ATP to AMP and pyrophosphate (PP_i_), which acts as a physiological “water softener” to control mineralization (Orriss et al., 2016). Additionally, ATP and its degradation product adenosine are themselves immunomodulatory molecules which can oppose or synergize with extracellular cGAMP. Since complete knockout or inhibition of ENPP1 disrupts essential physiology and alters immune signaling pathways, there is currently no tool to study the broad extracellular biology of cGAMP. Here, we developed an evolutionarily conserved substrate-selective mutation of ENPP1 to specifically manipulate extracellular cGAMP biology. We created a mouse strain harboring this mutation (*Enpp1^H362A^*) and found that these mice were more resistant to HSV-1 infection, suggesting a role for extracellular cGAMP in antiviral defense. Beyond HSV-1 infection, this cGAMP-selective ENPP1 mutation is a precise genetic tool to study the role of extracellular cGAMP in normal homeostasis, disease, and evolutionary history.

## Results

### Mutations of guanosine-adjacent residues in ENPP1 do not inhibit cGAMP hydrolysis

To design a mutagenesis scan for identifying substrate-selective ENPP1 mutations, we first compared the co-crystal structures of mouse ENPP1 bound to AMP, the product of ATP degradation (Kato et al., 2012), and mouse ENPP1 bound to pApG, the intermediate after the first phosphodiester bond cleavage of cGAMP (Kato et al., 2018). ENPP1 has a tight nucleotide-binding site, which the AMP portions of ATP and cGAMP occupy, and a secondary site, which only the guanine base of cGAMP occupies (guanosine-adjacent site) (**Fig. 1a-b**). Between the two sites, a cluster of residues chelate two zinc ions which position the α-phosphate of the substrates for nucleophilic attack by the catalytic residue T238.

**Fig. 1.**
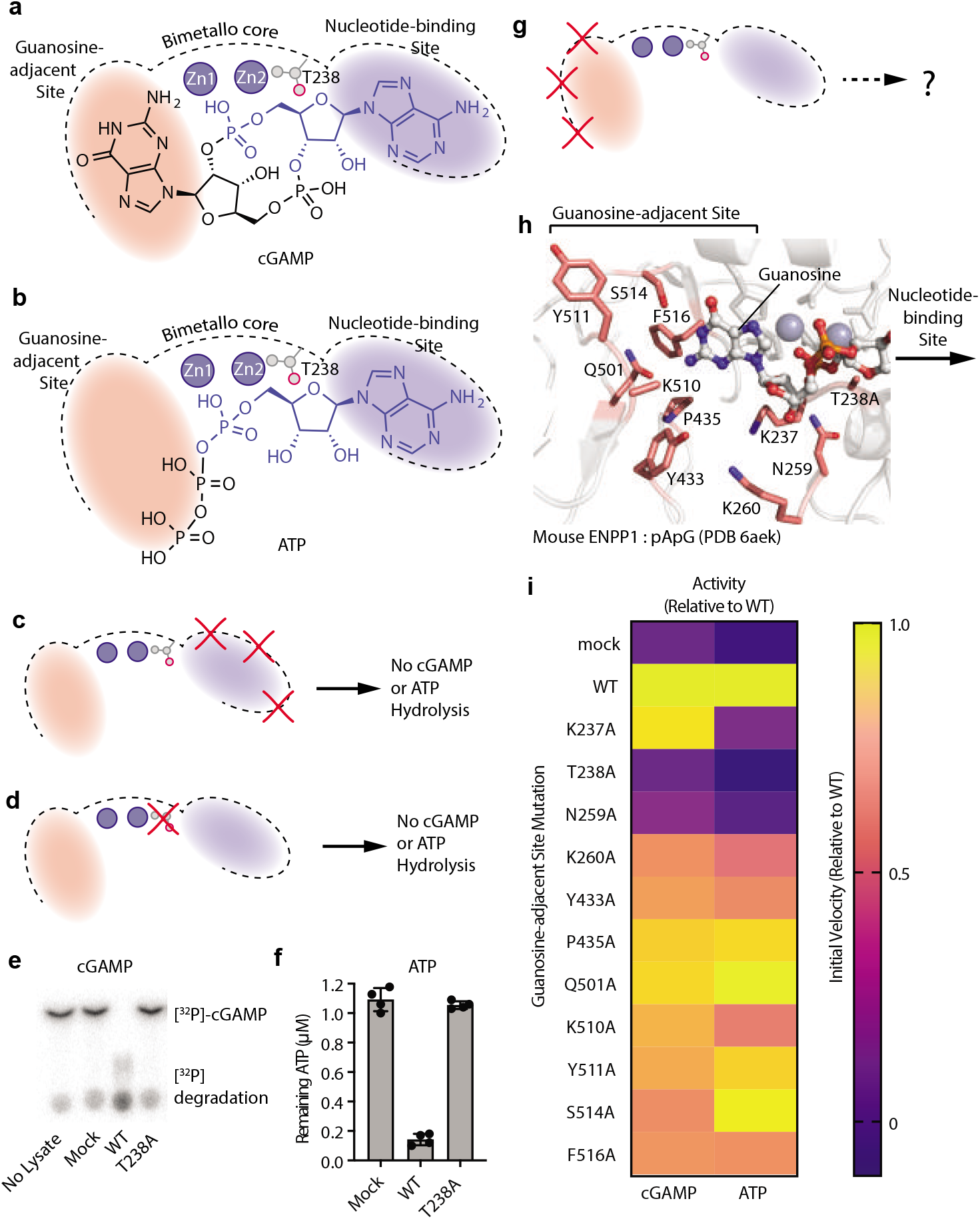
Mutations of guanosine-adjacent residues in ENPP1 do not inhibit cGAMP hydrolysis. **a-b** Schematic illustration of the ENPP1 active site bound to substrates cGAMP (**a**) and ATP (**b**). **c-d** Schematic illustration of mutations (red X’s) of the nucleotide-binding site (blue) (**c**) and nucleophile T238 (ball and stick) (**d**), leading to no substrate hydrolysis. **e** TLC cGAMP degradation assay for ENPP1 mutants expressed in cell lysate. Representative data shown from 3 independent reactions. **f** Luciferase-based ATP degradation assay for ENPP1 mutants expressed in cell lysate. *n* = 3 independent reactions, mean ± SD. **g** Schematic illustration of proposed mutations (red X’s) of the residues in the guanosine-adjacent site (red). **h** Mouse ENPP1 (gray cartoon) in complex with pApG (gray ball and sticks) (PDB:6aek). Guanosine-adjacent residues are colored salmon. **i** Heat map showing the initial velocity relative to WT of guanosine-adjacent residue mutations for the substrates cGAMP and ATP at pH 9. The mean initial velocity was calculated from a linear fit of the degradation reactions during early time points. *n* = 4 independent reactions.

We first mutated residues that we predicted might contribute to cGAMP binding without altering ATP binding. Previous studies showed that mutating key nucleotide-binding site residues inhibits ATP hydrolysis (Kato et al., 2012) (**Fig. 1c**), and we previously demonstrated that blocking the nucleotide-binding site with small molecule inhibitors blocks cGAMP hydrolysis (Carozza et al., 2020b). Therefore, we turned our attention to residues adjacent to the guanosine portion of cGAMP. First, we validated our assays using ENPP1^WT^ and the nucleophile mutation ENPP1^T238A^, which should have no hydrolysis activity toward either substrate (**Fig. 1d**). We generated whole cell lysates by overexpressing FLAG-tagged ENPP1 variants in 293T *ENPP1*^-/-^ cells (**Supplementary Fig. 1a**) as a fast and robust source of enzyme. [^32^P]-cGAMP degradation was monitored using thin layer chromatography (TLC) separation and autoradiography, and ATP degradation was monitored using a luciferase-based assay. Using these assays, we confirmed that ENPP1^WT^ is able to degrade both ATP and cGAMP, and ENPP1^T238A^ is not (**Fig. 1e-f**).

We then individually mutated ten residues with sidechains within 5 Å of the guanine base of cGAMP (**Fig. 1g–h; Supplementary Fig. 1b**) and measured the initial rate of substrate hydrolysis at both pH 9, the optimum pH for ENPP1 hydrolysis, and pH 7.5, the physiological pH (**Fig. 1i; Supplementary Fig. 1c-m**). Surprisingly, most of these mutations had only a modest effect on cGAMP degradation. ATP degradation activity usually tracked with cGAMP activity, suggesting that these mutations led to broad destabilizations as opposed to substrate-selective inhibition. Mutation of N259, a highly conserved residue that forms a hydrogen bond with the non-bridging phosphoryl oxygen of the substrates (Hausmann et al., 2011; Zalatan et al., 2006), inhibited activity for both substrates. The K237A mutation achieved the opposite of our goal by abolishing ATP degradation but preserving cGAMP degradation. It is possible that K237 is important for stabilizing the β- and □- phosphates of ATP, but too far away from the phosphodiester of cGAMP. Our mutagenesis scan revealed that it is not possible to selectively inhibit cGAMP hydrolysis by mutating guanosine-adjacent residues (**Fig. 1i**), and therefore an alternative strategy was required.

### Discovery and characterization of ENPP1^H362A^, a mutation that degrades ATP but not cGAMP

We next made alanine mutations of the six aspartate and histidine zinc-binding residues that comprise the catalytic core of ENPP1 (**Fig. 2a–b; Supplementary Fig. 2a**) and assessed their rates of substrate hydrolysis. All of the mutants were inactive toward both substrates except for the H362A mutation (ENPP1^H362A^); remarkably, this mutation recapitulated ENPP1^WT^ activity for ATP but had no detectable activity toward cGAMP (**Fig. 2c; Supplementary Fig. 2b–j**). We then mutated H362 to the other 18 amino acids, all of which were inactive toward cGAMP, indicating that histidine is essential at that position for cGAMP degradation (**Fig. 2d; Supplementary Fig. 3a–e**). Interestingly, a wide variety of amino acids could substitute for histidine at this position and still degrade ATP (**Fig. 2d**). Notably, lysine, alanine, glycine, threonine, and phenylalanine all had at least 90% activity compared to ENPP1^WT^. Additionally, charge-preserving mutations (e.g., histidine to lysine) generally preserved activity better than charge-flipping mutations (e.g., histidine to glutamate or aspartate). Finally, the purified ENPP1^H362A^ enzyme also has no cGAMP hydrolysis activity (**Fig. 2e**) but has ATP hydrolysis activity identical to ENPP1^WT^ (**Fig. 2f**). As expected, ENPP1^H362A^ was fully active toward the other NTP substrates of ENPP1 (GTP, UTP, and CTP) (**Fig. 2g; Supplementary Fig. 3f**), given the similarity between the binding postures of ATP and the other NTPs (Kato et al., 2012).

**Fig. 2.**
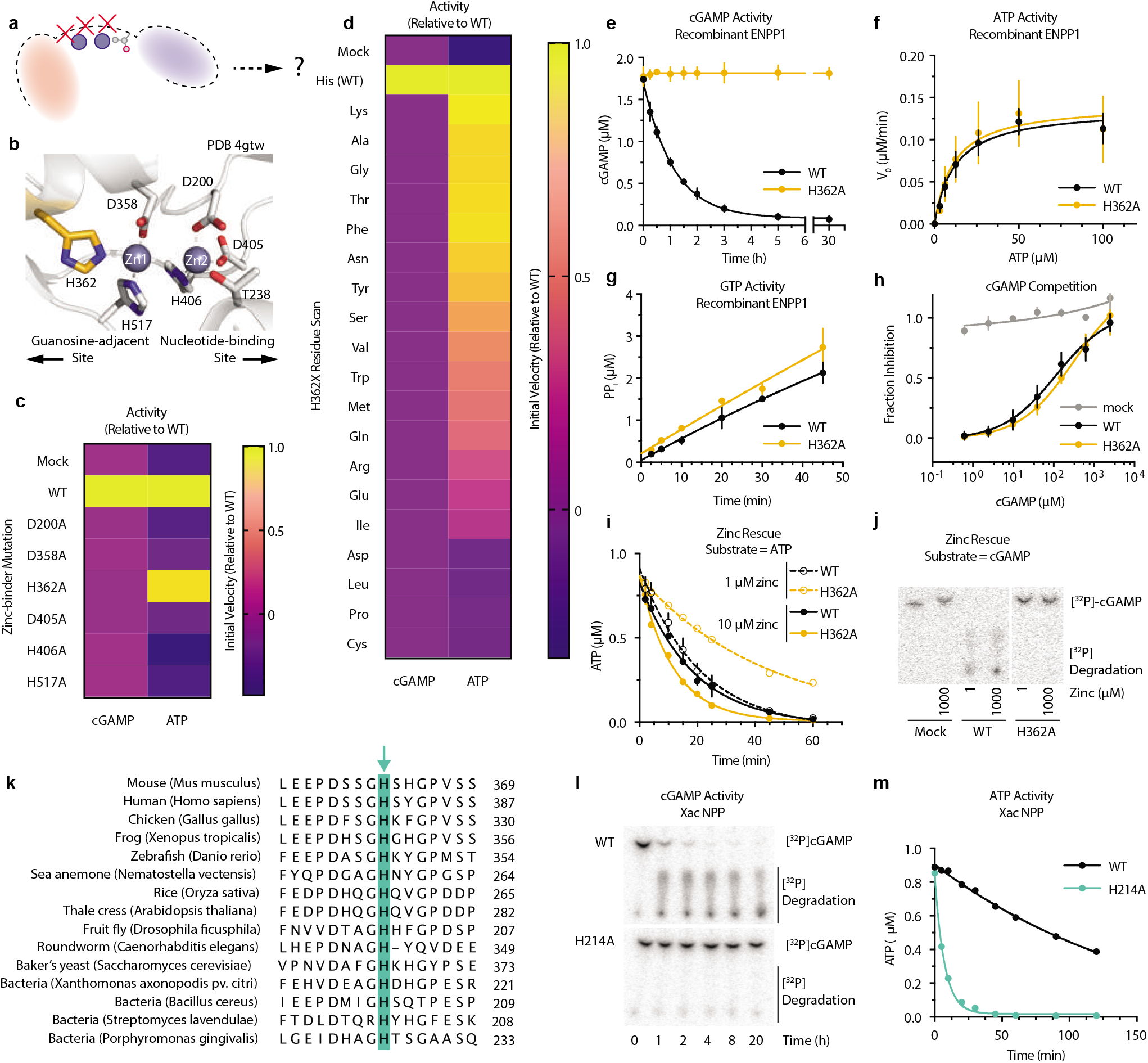
Discovery and characterization of ENPP1^H362A^, a mutation that degrades ATP but not cGAMP. **a** Schematic illustration of proposed mutations (red X’s) of the zinc-binding residues (zincs shown as dark gray spheres). **b** The catalytic center of mouse ENPP1 (PDB:4gtw). Zinc-binding residues shown as gray (D200, T238, D358, D405, H406, and H517) or orange (H362) sticks; zincs shown as dark gray spheres. **c** Heat map showing the initial velocity relative to WT of zinc-binding residues for the substrates cGAMP and ATP at pH 9. The mean initial velocity was calculated from a linear fit of the degradation reactions during early time points. *n* = 3 independent reactions. **d** Heat map showing the initial velocity relative to WT of H362X mutations (where X is each amino acid), for the substrates cGAMP and ATP at pH 9. The mean initial velocity was calculated from a linear fit of the degradation reactions during early time points. *n* = 3 independent reactions. **e** Kinetic analysis of cGAMP activity monitoring degradation products by TLC using recombinant purified ENPP1. *n* = 3 independent reactions, mean ± SD with some error bars too small to visualize. **f** Michaelis-Menten plots of ATP activity monitoring AMP production using recombinant purified ENPP1. *n* = 3 independent reactions, mean ± SEM. WT: *K_m_* = 12.1 μM, *k*_cat_ = 0.76 s^-1^, *k*_cat_/*K*_m_ = 6.3 x 10^4^ M^-1^s^-1^. H362A: *K*_m_ = 11.1 μM, *k*_cat_ = 0.79 s^-1^, *k*_cat_/*K*_m_ = 7.1 x 10^4^ M^-1^s^-1^. **g** Kinetic analysis of GTP monitoring pyrophosphate production using recombinant purified ENPP1. *n* = 2 independent reactions, mean ± SD. **h** Competitive inhibition of ATP hydrolysis with cGAMP at pH 7.5. *n* = 3 independent reactions, mean ± SD. WT: *K*_i_ = 120 μM. H362A: *K*_i_ = 320 μM. **i** Zinc rescue experiment with ATP as the substrate. *n* = 3 independent reactions, mean ± SD. **j** Zinc rescue experiment with cGAMP as the substrate. Data is representative of 3 independent reactions. **k** Sequence alignment of NPP from a range of species showing the conserved histidine (highlighted in green box). **l-m** TLC of cGAMP (**l**) and ATP (**m**) degradation by *Xac* NPP^WT^ compared to *Xac* NPP^H214A^ (both at 3 μM enzyme concentration). l representative of 3 independent reactions from 2 independent protein purifications. m *n* = 3 independent reactions, mean ± SD with some error bars too small to visualize.

We then investigated the mechanism of substrate selectivity of the ENPP1^H362A^ mutant. We first determined if ENPP1^H362A^ could still bind to cGAMP by testing whether cGAMP is a competitive inhibitor to the ENPP1-ATP reaction. ENPP1^WT^ and ENPP1^H362A^ had very similar *K*_i_ values, demonstrating that ENPP1^H362A^ could bind cGAMP equally well as ENPP1^WT^ and that binding affinity is driven by the tight interactions in the nucleotide-binding pocket (**Fig. 2h**). Since we did not observe any intermediates in the cGAMP TLC assay (**Supplementary Fig. 3g**), it is unlikely that ENPP1^H362A^ hydrolyzes the 2’-5’ phosphodiester bond but is unable to release the linear intermediate. We next tested whether ENPP1^H362A^ can hydrolyze cGAMP when given higher zinc concentrations, as H362 is also the right distance to chelate zinc. Interestingly, while increasing the zinc concentration from 1 μM to 10 μM sped up the rate of ATP hydrolysis for ENPP1^H362A^ with no effect on ENPP1^WT^ (**Fig. 2i**), we were unable to rescue cGAMP activity even with 1 mM zinc, which is 100-fold above physiological conditions (10-20 μM) (**Fig. 2j**). These results demonstrate that although H362A has an impact on zinc binding at sub-physiological concentrations of zinc, this slight deficiency is not responsible for the loss of cGAMP activity. The crystal structure of ENPP1 with pApG (Kato et al., 2018) shows that H362 and the guanine ring of cGAMP are ~4.5 Å apart and aligned parallel to one another in a stacking interaction. Furthermore, ENPP1 is unable to degrade but still binds 3’3’-cGAMP (Li et al., 2014) due to inappropriate geometry to initiate the nucleophilic attack (Kato et al., 2018). Based on these findings, we hypothesize that H362 is necessary to π-π stack with the guanine ring of cGAMP so that the phosphodiester bond is aligned in the optimal geometry required for in-line nucleophilic attack.

Since ENPP1 is part of a highly conserved NPP protein family, we investigated the evolutionary conservation of this residue and its requirement for cGAMP degradation. This histidine is conserved from humans and mice through lower animals, plants, and bacteria (**Fig. 2k**). Previous studies had characterized NPP from one particular bacteria species, *X. axonopodis pv. citri* (*Xac*) (Sunden et al., 2017; Zalatan et al., 2006), although its role in cGAMP hydrolysis is unexplored. *Xac* NPP can degrade cGAMP, and the H214A mutation, which corresponds to the murine H362A mutation, similarly abolished its cGAMP hydrolysis activity (**Fig. 2l; Supplementary Fig. 3h**). Its ATP hydrolysis, however, was not only preserved, but also several-fold faster than WT (**Fig. 2m**). Furthermore, all of the eukaryotic sequences and all but one prokaryotic sequence used in the sequence alignment had a predicted signal peptide (**Supplementary Fig. 3i**), suggesting extracellular localization. The remarkable conservation of the corresponding histidine mutations in distinguishing between cGAMP and NPP’s other substrates, as well as the conserved extracellular localization, suggest that this mutant can be introduced into a wide variety of eukaryotic and prokaryotic organisms to study the pathophysiology of extracellular cGAMP signaling.

### *Enpp1^H362A^* mice can degrade ATP but not cGAMP

We next created homozygous *Enpp1^H362A^* mice using standard CRISPR-based homologous recombination (**Supplementary Fig. 4a–b**). After verifying the genotype of the mice (**Supplementary Fig. 4c**), we sought to confirm that these mice exhibited the expected enzymatic activity: intact ATP hydrolysis but defective cGAMP hydrolysis. We compared tissue lysates from *Enpp1^WT^* and *Enpp1^H362A^* mice to those from *Enpp1^asj^* mice, which harbor a point mutation that abolishes ENPP1 activity (Li et al., 2013a). While tissue lysates from *Enpp1^WT^* mice rapidly degraded [^32^P]-cGAMP, tissue lysates from *Enpp1^H362A^* mice did not, mirroring the deficiency seen in *Enpp1^asj^* mice (**Fig. 3a**). In addition to the ENPP1 tethered to the surface of cells, ENPP1 is also secreted into the circulation, allowing cGAMP degradation to occur in the plasma (Belli et al., 1993; Jansen et al., 2012). Plasma from *Enpp1^WT^* mice readily degraded cGAMP, while plasma from *Enpp1^H362A^* mice did not (**Fig. 3b**). Despite the inability to degrade cGAMP, basal tissue cGAMP levels were generally low and not significantly different between *Enpp1^WT^, Enpp1^H362A^*, and *Enpp1^asj^* mice (**Supplementary Fig. 4d**), suggesting that cGAMP is not exported in the absence of stimuli. Finally, we assessed cGAMP degradation *in vivo* by subcutaneously injecting cGAMP into each mouse, collecting plasma after 30 minutes, and measuring cGAMP concentration by mass spectrometry. cGAMP levels in the plasma of *Enpp1^H362A^* and *Enpp1^asj^* mice were >100-fold higher than in *Enpp1^WT^* mice (**Fig. 3c**), demonstrating that cGAMP hydrolysis is severely impaired in *Enpp1^H362A^* mouse plasma.

**Fig. 3.**
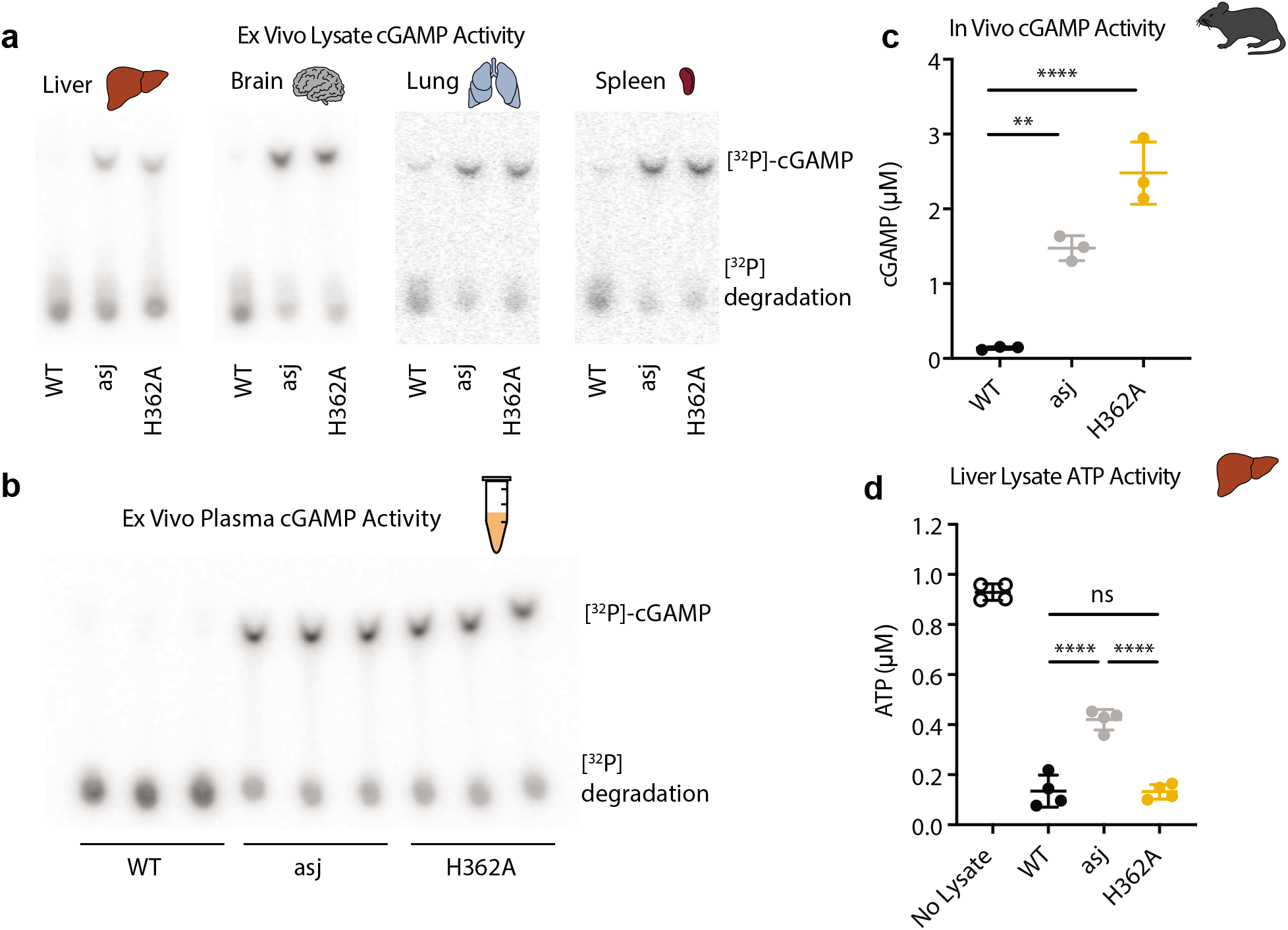
*Enpp1^H362A^* mice can degrade ATP but not cGAMP. **a** *Ex vivo* organ lysate (1 mg/mL) cGAMP degradation at pH 9 assessed by TLC after 4 hours (liver, brain) or 2 hours (lung, spleen). *n* = 1 mouse per genotype. **b** *Ex vivo* plasma cGAMP degradation assessed by TLC after 4 hours. *n* = 3 mice per genotype. **c** *In vivo* cGAMP degradation. 5 mg/kg of cGAMP was injected subcutaneously into each mouse. Blood was collected from each mouse after 30 minutes and cGAMP was measured by LC-MS/MS. *n* = 3 mice per genotype. **d** *Ex vivo* liver lysate (1 mg/mL) ATP degradation at pH 7.5 assessed after 20 minutes. *n* = 4 mice per genotype. For **c-d** data are shown as the mean ± SD. *p* values were calculated by unpaired *t* test with Welch’s correction. \*\**p* < 0.01, \*\*\*\**p* < 0.0001.

We then assessed ATP activity in these mice using liver lysates. The liver was the only organ suitable for these assays, as it expresses high levels of ENPP1, while ENPP1-independent ATP degradation predominates in other organs, precluding them from being used in this assay. Liver lysates from *Enpp1^asj^* mice had a marked deficiency in ATP degradation, but lysates from *Enpp1^H362A^* mice degraded ATP as well as *Enpp1^WT^* liver lysates, confirming that tissue ENPP1^H362A^ retains the ability to degrade ATP (**Fig. 3d; Supplementary Fig. 4e**). Taken together, *Enpp1^H362A^* mice are unable to degrade cGAMP but retain the ability to degrade ATP.

### *Enpp1^H362A^* mice do not exhibit abnormal calcification despite low plasma pyrophosphate

*Enpp1^asj^* mice exhibit progressive calcification of joints, vasculature, and soft tissue, resulting in premature death (Li et al., 2013a). An analogous disease, known as generalized arterial calcification of infancy (GACI), occurs in humans with inactivating ENPP1 mutations. As with the *Enpp1^asj^* mice, GACI patients present with widespread arterial calcification and death in early infancy (Rutsch et al., 2001, 2003, 2008). It is hypothesized that the aberrant calcification found in ENPP1-deficient mice and humans is due to the inability to degrade extracellular ATP, leading to a deficiency in PP_i_, which is known to regulate mineralization (Rutsch et al., 2008). However, since cGAMP was only recently identified as a substrate of ENPP1, it is unknown if impaired cGAMP metabolism also leads to the aberrant calcification observed in humans and mice.

As cGAMP degradation is uncoupled from extracellular ATP degradation in *Enpp1^H362A^* mice, they are an ideal model for determining the role of extracellular cGAMP in the progressive calcification seen in *Enpp1^asj^* mice. *Enpp1^H362A^* mice have a normal lifespan and readily breed, unlike the *Enpp1^asj^* mice, which have difficulty breeding beyond 2–3 months of age due to worsening arthritis. The *Enpp1^H362A^* mice do not exhibit the gross joint calcification seen in *Enpp1^asj^* mice (**Fig. 4a**). In contrast to tissues from *Enpp1^asj^* mice, histological analysis of tissues from 20-week-old *Enpp1^H362A^* mice showed very minimal calcification (**Fig. 4b**). Taken together, these results suggest that extracellular cGAMP does not play a significant role in the calcification seen in *Enpp1^asj^* mice, and that this phenotype is likely due to the disruption of extracellular ATP metabolism.

**Fig. 4.**
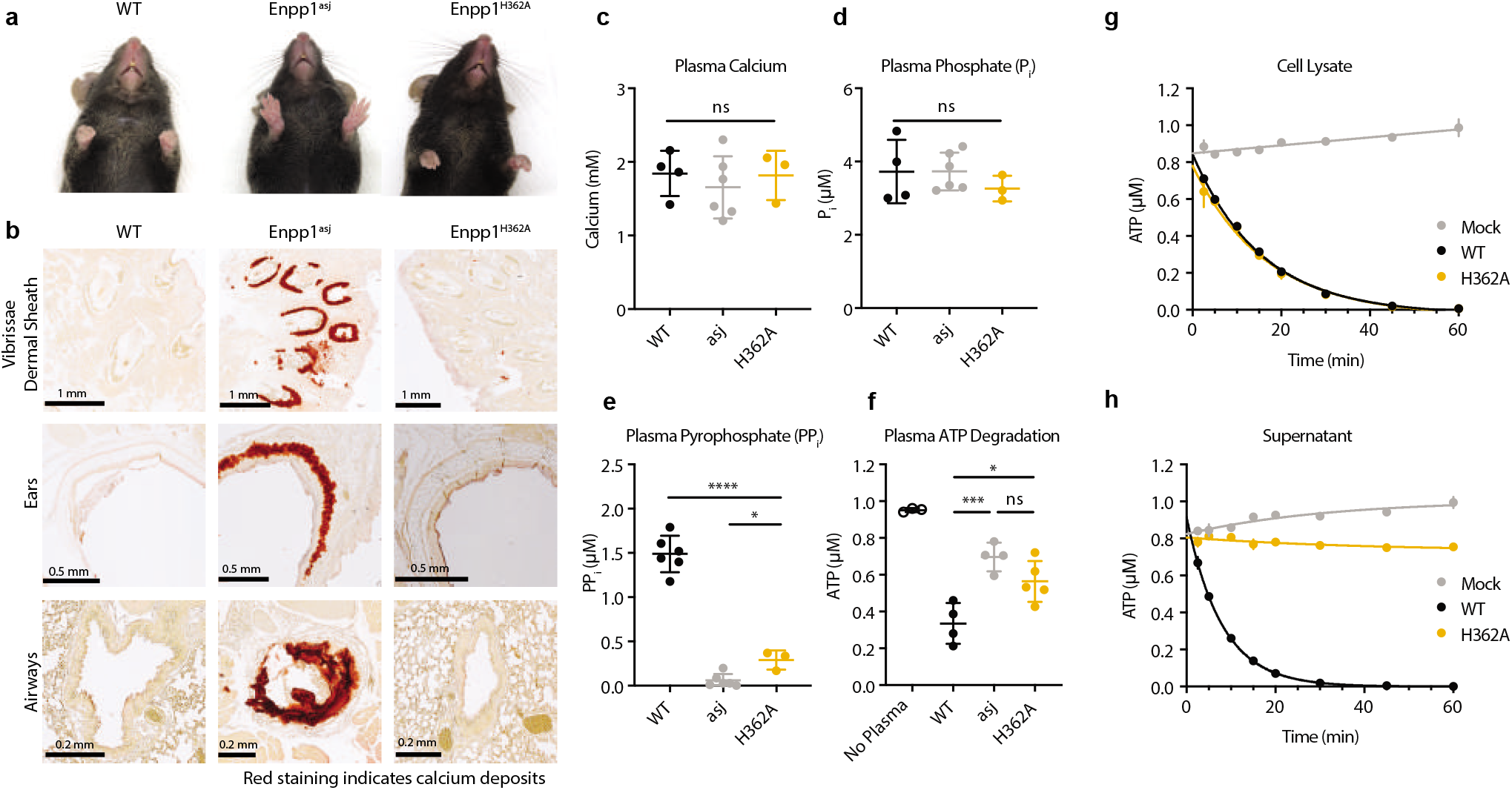
*Enpp1^H362A^* mice do not exhibit systemic calcification despite low plasma pyrophosphate. **a** Photos of WT, *Enpp1^H362A^*, and *Enpp1^asj^* mouse paws. The *Enpp1^asj^* mouse paws are unable to relax due to the calcified joints. **b** 20-week-old mice were euthanized, and the ears, airways, and vibrissae were fixed and stained with alizarin red to identify calcium deposits. Representative images are shown from 1–2 mice per genotype. **c-d** Plasma calcium (**c**) and phosphate (**d**) concentrations in *Enpp1^WT^, Enpp1^asj^*, and *Enpp1^H362A^* mice. *n* = 4 *Enpp1^WT^*, 6 *Enpp1^asj^*, and 3 *Enpp1^H362A^* mice. **e** Plasma pyrophosphate concentrations in concentrations in *Enpp1^WT^, Enpp1^asj^*, and *Enpp1^H362A^* mice. *n* = 6 *Enpp1^WT^*, 6 *Enpp1^asj^*, and 3 *Enpp1^H362A^* mice. **f** *Ex vivo* plasma ATP degradation at pH 7.5 assessed by luciferase assay after 45 minutes in *Enpp1^WT^, Enpp1^asj^*, and *Enpp1^H362A^* mice. *n* = 4 *Enpp1^WT^*, 4 *Enpp1^asj^*, and 5 *Enpp1^H362A^* mice. **g-h** *In vitro* ATP degradation comparing overexpressed ENPP1^WT^ and ENPP1^H362A^ as cell-surface protein from cell lysate (**g**) and as secreted protein from cell supernatant (**h**). For **c-f** data are shown as the mean ± SD. *p* values were calculated by unpaired *t* test with Welch’s correction. \**p* < 0.05, \*\*\**p* < 0.001, \*\*\*\**p* < 0.0001.

*Enpp1^asj^* mice are reported to have normal plasma calcium and phosphorous levels, while their plasma PPi levels are significantly lower than normal due to their inability to degrade ATP (Li et al., 2013a). Like *Enpp1^WT^* and *Enpp1^asj^* mice, *Enpp1^H362A^* mice had normal plasma calcium and phosphorous levels (**Fig. 4c–d**). However, while *Enpp1^WT^* mice had a normal level of plasma PP_i_ (1.5 μM), *Enpp1^H362A^* and *Enpp1^asj^* mice had 300 nM and 60 nM plasma PP_i_, respectively (**Fig. 4e**). This was an unexpected finding, as *Enpp1^H362A^* liver lysate did not exhibit a defect in ATP degradation (**Fig. 3d**). This raised the possibility that plasma ENPP1^H362A^ does not exhibit the same behavior as tissue ENPP1^H362A^. Indeed, *Enpp1^H362A^* mouse plasma was defective in ATP degradation, similar to *Enpp1^asj^* (**Fig. 4f, Supplementary Fig. 5a**). Our *in vitro* experiments suggested that ATP hydrolysis by ENPP1^H362A^ is partly sensitive to zinc concentration. However, supplementing the reaction with 100 μM zinc did not restore plasma ATP activity (**Supplementary Fig. 5b**). Nevertheless, our results suggest that the low plasma PP_i_ concentration does not contribute to the calcification phenotype in *Enpp1^asj^* mice.

We hypothesized that the difference between plasma and tissue ENPP1^H362A^ activity was due to differences between the secreted ENPP1 found in plasma and the cell-surface ENPP1 found in tissues. To test this hypothesis, we used our *in vitro* activity assay to compare secreted ENPP1 in cell supernatants with cell-surface ENPP1 present in cell lysates. Unlike the cell-surface ENPP1^H362A^, the secreted ENPP1^H362A^ was defective in ATP degradation (**Fig. 4g–h; Supplementary Fig. 5c–d**). This discrepancy was not due to a difference in ENPP1 secretion since ENPP1^WT^ and ENPP1^H362A^ were present at similar amounts in the supernatant (**Supplementary Fig. 5e**). Therefore, it is likely that the soluble ENPP1^H362A^ in mouse plasma is unable to degrade ATP, while cell-surface ENPP1^H362A^ in mouse tissue retains the ability to degrade ATP. Although we currently lack the ability to measure extracellular PPi levels in tissues, we surmise that tissue PP_i_ is normal in *Enpp1^H362A^* mice and that tissue PP_i_, rather than plasma PP_i_, is important for preventing tissue calcification.

### *Enpp1^H362A^* mice are resistant to HSV-1 infection

Although extracellular cGAMP does not play a role in ENPP1-mediated systemic calcification, there are many contexts in which extracellular cGAMP likely plays an important role as an immunotransmitter. A previous study found that mice lacking the cGAMP importer LRRC8E had a diminished immune response and higher viral titers in response to herpes simplex virus 1 (HSV-1) infection (Zhou et al., 2020a). However, LRRC8E has another role as part of the volume-regulated ion channel (VRAC), a critical regulator of cell size which transports a wide variety of substrates (Qiu et al., 2014; Voss et al., 2014). As ENPP1^H362A^ is an ideal tool to specifically probe extracellular cGAMP biology, we were able to formally determine if extracellular cGAMP was responsible for the observed resistance to HSV-1 infection. We systemically infected *Enpp1^WT^, Enpp1^H362A^*, and *Enpp1^asj^* mice with HSV-1. Compared to WT mice, *Enpp1^H362A^* mice had significantly lower levels of the viral gene *HSV-gB* in the liver, lung, and spleen (**Fig. 5a**). This was accompanied by lower viral loads in *Enpp1^H362A^* spleen and kidney lysates (**Supplementary Fig. 6a**). Furthermore, *Enpp1^H362A^* mice exhibited significantly decreased expression of *Ifnb1* and the downstream cytokines *Cxcl10* and *Il6* in the liver, lung, and spleen, as well as decreased *Tnfa* in the liver (**Fig. 5b–e**); this decrease in cytokine production was likely due to lower viral loads in the mice at this time point. Taken together, these data indicate that *Enpp1^H362A^* mice are significantly more resistant to HSV-1 infection, suggesting that enhanced extracellular cGAMP is protective against viral infection.

**Fig. 5.**
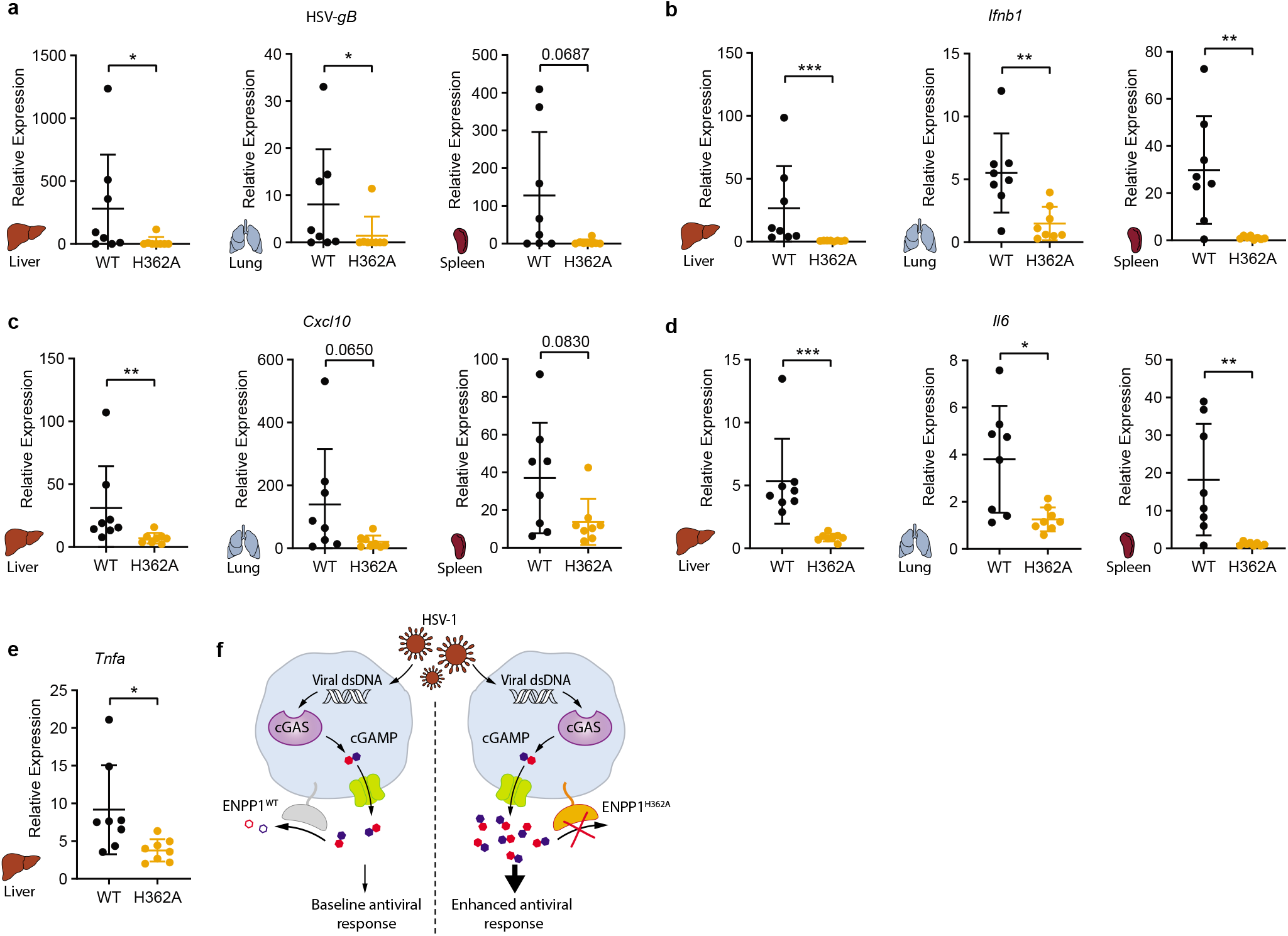
*Enpp1^H362A^* mice are resistant to HSV-1 infection. **a-e** Mice were inoculated with 2.5 x 10^7^ PFU/mouse HSV-1 through intravenous injection. The mice were euthanized after 12 h, and RNA was isolated from the liver, lungs, and spleen. RT-qPCR was performed to determine the expression levels of *HSV-gB* (**a**), *Ifnb1* (**b**), *Cxcl10* (**c**), *Il6* (**d**), and *Tnfa* (**e**). Transcript levels were normalized to the average of 2 uninfected mice per genotype. n = 8 infected mice per genotype. Data are shown as the mean ± SD. p values were calculated using the Mann-Whitney test. \**p* < 0.05, \*\**p* < 0.01, \*\*\**p* < 0.001; p value is shown if between 0.05 and 0.1. **f** Model of cGAMP as a paracrine immunotransmitter regulated by ENPP1 during viral infection. Infected cells produce and export cGAMP in response to viral dsDNA. When ENPP1^WT^ is present, extracellular cGAMP is degraded, diminishing the baseline antiviral response. When cGAMP-specific ENPP1^H362A^ is present, extracellular cGAMP accumulates, leading to an enhanced antiviral response.

The response to HSV-1 infection in *Enpp1^asj^* mice largely mirrors the response in *Enpp1^H362A^* mice, although in some cases there was no difference between *Enpp1^WT^* mice and *Enpp1^asj^* mice (**Supplementary Fig. 6b–g**). For example, while there was a highly significant decrease in splenic *Ifnb1* in *Enpp1^H362A^* mice (**Fig. 5b**), there was no discernable difference between *Enpp1^WT^* and *Enpp1^asj^* mice (**Supplementary Fig. 6d**). It is possible that the impaired ATP metabolism or the pathophysiology and overall poor health of *Enpp1*^asj^ mice counteracts the antiviral effect of extracellular cGAMP, further highlighting the importance of *Enpp1^H362A^* mice in dissecting extracellular cGAMP biology. Interestingly, we did not detect cGAMP in the plasma of any of the mice, with an assay detection limit of 85 pg/mL (125 pM) (**Supplementary Fig. 6h**). This suggests that plasma cGAMP levels are at least 10- to 100-fold lower than cGAMP’s *K*_d_ toward STING and are therefore unlikely to be physiologically relevant. The lack of circulating cGAMP suggests that extracellular cGAMP is acting locally as a paracrine immunotransmitter to confer resistance to HSV-1 infection (**Fig. 5f**).

## Discussion

In this study we identified and characterized ENPP1^H362A^, a single amino acid mutation in ENPP1 that abolishes cGAMP degradation but leaves ATP degradation intact. Our active site mutagenesis scan also discovered a mutation with the opposite substrate selectivity: ENPP1^K237A^ abolishes ATP degradation but preserves cGAMP degradation. While this report used the H362A mutation to study extracellular cGAMP-specific biology, the ATP-selective K237A mutation could be used to study extracellular ATP-specific biology in the future. These discoveries add to the small but growing field of identifying substrate-selective enzyme mutations (McKinney and Cravatt, 2006) and substrate-selective enzyme inhibitors (Maianti et al., 2019). This technique is useful for studying both the mechanisms of substrate catalysis and the physiology of specific substrates.

It was particularly surprising that mutating H362 to a variety of amino acids still preserved ATP activity, given that this histidine seems to be chelating one of the two essential zinc atoms in the crystal structures (Dennis et al., 2020; Jansen et al., 2012; Kato et al., 2012, 2018). Additionally, this histidine is highly conserved in members of the NPP and alkaline phosphatase superfamily, which extends from mammals to bacteria (Gijsbers et al., 2001; Zalatan et al., 2006). Interestingly, our data suggest that ENPP1^H362A^ is still able to bind zinc to hydrolyze ATP, likely with a slight decrease in affinity. It is puzzling why H362 is so conserved if it is completely unnecessary for NTP hydrolysis. It is possible that histidine becomes more favorable in zinc-poor environments, or that it has other destabilizing effects that we did not identify in our experiments. Since cGAMP-STING signaling is conserved in invertebrates, it is tempting to hypothesize that extracellular cGAMP biology is evolutionarily ancient and that NPP-mediated cGAMP degradation is a key negative regulatory mechanism. We propose that this mutant can be introduced into many organisms to study the origin of extracellular cGAMP pathophysiology, opening another area of research in this nascent field.

To further demonstrate the utility of this mutation, we created *Enpp1^H362A^* mice to study the physiological roles of extracellular cGAMP metabolism *in vivo*. Unlike *Enpp1*^asj^ mice, *Enpp1^H362A^* mice do not exhibit widespread calcification or early death, emphasizing that this phenotype is due to impaired ATP hydrolysis, rather than an increase in extracellular cGAMP. Interestingly, *Enpp1^H362A^* mice have significantly decreased plasma PPi levels, despite not having any evidence of disease, suggesting that low plasma PP_i_ does not actually play a role in the pathophysiology of tissue calcification. Instead, local tissue PP_i_ may be more important in preventing calcification. Supporting this hypothesis, a recent study found that osteoblast-specific loss of ENPP1 led to increased bone volume, decreased trabecular spacing, and increased matrix calcification, without any change to plasma PP_i_ (Roberts et al., 2020).

The *Enpp1^H362A^* mouse model is a robust tool that can be used for a wide range of precise genetic experiments to study extracellular cGAMP in health and disease. Using *Enpp1^asj^* mice alters both the extracellular cGAMP and the extracellular ATP concentrations, and these mice suffer from severe health complications and premature death. While the *Enpp1^H362A^* mice largely resemble WT mice at baseline, they diverge in their response to viral infection. *Enpp1^H362A^* mice are significantly more resistant to HSV-1 infection, indicating that extracellular cGAMP is protective against this virus. Interestingly, the lower viral load in these mice was accompanied by lower levels of IFN-β and other cytokines. This may simply be a consequence of the timing of this assay or may indicate that increased STING signaling led to a paradoxical decrease in cytokine production, as previously reported for STING-mediated responses to RNA viruses (Franz et al., 2018). With its demonstrated utility in studying the response to HSV-1, the *Enpp1^H362A^* mouse model will revolutionize the extracellular cGAMP field, finally enabling precise genetic experiments to study extracellular cGAMP and its regulation by ENPP1 in a wide range of disease contexts. As we expand our knowledge of the importance of extracellular cGAMP signaling to new diseases and physiological settings, the ability to regulate extracellular cGAMP will provide new therapeutic opportunities to manipulate and treat these diseases.

## Methods

### Synthesis and purification of cGAMP and [^32^P]-cGAMP

To enzymatically synthesize cGAMP (Ritchie et al., 2019), 1 μM purified sscGAS was incubated with 20 mM Tris-HCl pH 7.4, 2 mM ATP, 2 mM GTP, 20 mM MgCl_2_, and 100 µg/mL herring testis DNA (Sigma) for 24 h. The reaction was then heated at 95°C for 3 min and filtered through a 3-kDa filter. cGAMP was purified from the reaction mixture using a PLRP-S polymeric reversed phase preparatory column (100 Å, 8 μm, 300 x 25 mm; Agilent Technologies) on a preparatory HPLC (1260 Infinity LC system; Agilent Technologies) connected to UV-vis detector (ProStar; Agilent Technologies) and fraction collector (440-LC; Agilent Technologies). The flow rate was set to 25 mL/min. The mobile phase consisted of 10 mM triethylammonium acetate in water and acetonitrile. The mobile phase started as 2% acetonitrile for the first 5 min. Acetonitrile was then ramped up to 30% from 5-20 min, then to 90% from 20-22 min, maintained at 90% from 22-25 min, and then ramped down to 2% from 25-28 min. Fractions containing cGAMP were lyophilized and resuspended in water. The concentration was determined by measuring absorbance at 280 nm. To enzymatically synthesize [^32^P]-cGAMP, 1 μM purified sscGAS was incubated with 20 mM Tris-HCl pH 7.4, 250 µCi (3000 Ci/mmol) [α-^32^P]-ATP (Perkin Elmer), 1 mM GTP, 20 mM MgCl_2_, and 100 µg/mL herring testis DNA (Sigma) in a reaction volume of 100 μL for 24 h. The reaction was purified by preparatory TLC on a HP-TLC silica gel plate (Millipore), eluted in water, and filtered through a 3-kDa filter to remove silica gel.

### Cell culture

HEK 293T *ENPP1*^-/-^ cells were generated in a previous study (Carozza et al., 2020a). Vero and Expi293F cell lines were gifts from Dr. Peter Kim (Stanford University). 293T *ENPP1*^-/-^ and Vero cell lines were maintained in a 5% CO_2_ incubator at 37 °C in DMEM (Corning Cellgro) supplemented with 10% FBS (Atlanta Biologics) and 100 U mL^-1^ penicillin-streptomycin (Thermo Fisher). The Expi293F cell line was maintained in an 8% CO_2_ incubator at 37 °C shaking in baffled flasks, in a mixture of 66% FreeStyle Expression Media (Thermo) and 33% Expi293 Expression Media (Thermo). All cell lines tested negative for mycoplasma contamination.

### Recombinant DNA

The pcDNA3-mouseENPP1-FLAG plasmid was synthesized by Genscript. The *Xac* NPP-MBP-pMAL plasmid was a gift from Dr. Daniel Herschlag (Sunden et al., 2017). The single point mutations were introduced using QuikChange mutagenesis (Agilent) and verified by sequencing the region of the mutation. Primers used for mutagenesis are shown in **Supplementary Table 1**.

### cGAMP activity assays

cGAMP activity assays (20 μL total) were composed of the following: cell or organ lysate (50%) or recombinant ENPP1 (1 – 10 nM), cGAMP (1 to 5 μM, with trace [^32^P]-cGAMP spiked in), and buffer (standard assay buffer unless otherwise noted was 100 mM Tris pH 9 or pH 7.5, 150 mM NaCl, 500 μM CaCl_2_, 10 μM ZnCl_2_). At indicated times, 1 μL aliquots of the reaction were quenched by spotting on HP-TLC silica gel plates (Millipore). The TLC plates were run in mobile phase (85% ethanol, 5 mM NH_4_HCO_3_) and exposed to a phosphor screen (GE BAS-IP MS). Screens were imaged on a Typhoon 9400 scanner.

### ATP activity assays

ATP activity assays (10 μL total in a 384 well PCR plate) were composed of the following: cell or organ lysate (0.1–1%, depending on ENPP1 expression level) or recombinant ENPP1 (1 – 3 nM), 1 μM ATP (Sigma) and buffer (standard assay buffer unless otherwise noted was 100 mM Tris pH 9 or pH 7.6, 150 mM NaCl, 500 μM CaCl_2_, 10 μM ZnCl_2_). Reactions were started at indicated times and ended simultaneously by heating at 95 °C for 10 minutes. Reactions (5 μL) were transferred to a white 384 well plate, mixed with CellTiterGlo (5 μL), and luminescence was read after 15 minutes on a Tecan Spark plate reader.

### Additional enzyme assays

For the GTP, CTP, and UTP activity assays, generation of pyrophosphate was measured according to previously published method (Oheim et al., 2020). To convert pyrophosphate into ATP, 5 µL of each plasma sample was added to a reaction mixture consisting of 16 μM adenosine phosphosulfate, 80 mM MgSO_4_, 50 mM HEPEs, and 0.1 µL ATP sulfurylase (MCLab). The reaction mixture was then incubated at 37°C for 10 min, followed by 90°C for 10 min to inactivate the enzyme. In order to measure ATP, 25 μL of the reaction mixture was added to 25 μL of CellTiter-Glo (Promega). Luminescence was measured after 10 min using a 0.5 s integration time. For measurement of *k*_cat_/*K*_m_ for recombinant ENPP1 with the substrate ATP, the commercial AMPGlo kit (Promega) was used according to the manufacturer’s instructions.

### Recombinant mouse ENPP1 purification

Procedures for culturing and transfecting Expi293F cells (Thermo Fisher) were based on the manufacturer’s instructions. One day prior to transfection, the cells were split to 3×10^6^ cells/mL in baffled flasks (Corning). On the day of transfection, cells were diluted to 3-4 ×10^6^ cells/mL (if not already within the range) and transfected with plasmid DNA (0.5 μg DNA/mL cells) using FectoPro (Polyplus) (1 μL FectoPro/mL cells). Cells were immediately boosted with valproic acid (3 μM) and D-glucose (4 g/L). Cells were cultured for an additional 3 days. The media was harvested by centrifuging at 1000 x *g* for 10 minutes and passing through a 0.45 μm filter. Media was batch bound with HisPur cobalt resin (Thermo Fisher) (1 mL resin/ 60 mL culture) for 1 hour at 4 °C then loaded onto a fritted column. The column was washed two times with 2 column volumes (CV) wash buffer 1 (20 mM Tris pH 8.0, 150 mM NaCl, 10 mM imidazole) and once with 2 CV wash buffer 2 (20 mM Tris pH 8.0, 150 mM NaCl, 20 mM imidazole). Four elutions were performed with 1 CV elution buffer (20 mM Tris pH 8.0, 150 mM NaCl, 300 mM imidazole). Elution fractions were pooled and dialyzed against dialysis buffer (20 mM Tris pH 7.4, 150 mM NaCl) overnight at 4 °C. Protein was concentrated to 1 mg/mL in 20 mM Tris pH 7.4, 150 mM NaCl, 10% glycerol, and snap frozen for storage at −80 °C. Yield was ~1 mg ENPP1 per 60 mL of Expi293F culture.

### Recombinant *Xac* NPP-MBP purification

*Xac* NPP-MBP was expressed in BL21(DE3) cells. 1 L of bacteria were induced at OD 0.6 with 0.3 M IPTG and grown at 30 °C for 16 hours. Bacteria were pelleted at 4000 x g and resuspended in column buffer (50 mM Tris pH 8, 150 mM NaCl, 20 µM ZnCl_2_. After two freezethaw cycles in liquid nitrogen, lysate was sonicated, centrifuged at 40,000 x *g* for 45 minutes, and filtered through a 0.45 μm filter. Lysate was flowed over 1 mL of streptactin resin, washed with 20 CV of column buffer, and eluted with column buffer + 2.5 mM desthiobiotin. Fractions containing *Xac* NPP-MBP were pooled and dialyzed overnight against 50 mM Tris pH 8, 150 mM NaCl, 100 μM ZnCl_2_.

### NPP family sequence alignment and signal peptide prediction

Representative species were selected from classical model organisms and the EMBL-EBI protein family (Phosphodiest PF01663). Multiple sequences alignment was performed using MAFFT and visualized using Jalview. Protein accession numbers are as follows: Mouse (*Mus musculus*) P06802; Human (*Homo sapiens*) P22413; Chicken (*Gallus gallus*) XP_040523241.1; Frog (*Xenopus tropicalis*) XP_031758349.1; Zebrafish (*Danio rerio*) NP_001025339.1; Sea anemone (*Nematostella vectensis*) XP_032232033.1; Rice (*Oryza sativa*) XP_025879414.1; Thale cress (*Arabidopsis thaliana*) NP_194697.1; Roundworm (*Caenorhabditis elegans*) NP_001041086.1; Baker’s yeast (*Saccharomyces cerevisiae*) PTN13750.1; Bacteria (*Xanthomonas axonopodis pv. citri*) Q8PIS1_XANAC; Bacteria (*Bacillus subtilis*); Bacteria (*Streptomyces lavendulae*) ATZ26543.1; Bacteria (*Porphyromonas gingivalis*) PDP83833.1; Bacteria (*Bacillus cereus*) Q73E52_BACC1. Signal peptides were predicted using SignalP-5.0 and transmembrane domains were predicted using TMHMM (DTU Bioinformatics).

### Mouse models

C57BL/6J (Stock #000664), B6(C)-Cgas^tm1d(EUCOMM)Hmgu^/J (Stock # 026554), and C57BL/6J-*Enpp1^asj^*/GrsrJ (Stock #012810) mice were purchased from the Jackson Laboratory. Male and female mice were included in every experiment, unless otherwise noted. Mice were maintained at Stanford University in compliance with the Stanford University Institutional Animal Care and Use Committee (IACUC) regulations. All procedures were approved by the Stanford University Administrative Panel on Laboratory Animal Care (APLAC).

### Generating the transgenic *Enpp1^H362A^* mouse strain

First, single-guide RNAs (sgRNAs) were designed against the H362 locus in exon 9 of Enpp1 using publicly available design tools (Haeussler et al., 2016) (**Supplementary Table 1**). The sgRNA was then complexed with Alt-R S.p. Cas9 nuclease (Integrated DNA Technologies) as a ribonucleoprotein (RNP) particle. We then designed a donor sequence to serve as the template for homologous recombination (**Supplementary Table 1**). The 100 nucleotide-long donor sequence contained blocking mutations near the PAM sequence to prevent repeated editing (Okamoto et al., 2019). The donor sequence was then synthesized as single-stranded DNA (Integrated DNA Technologies). The RNP particles and donor template were microinjected into the pronuclei of one-cell embryos from C57BL/6 mice, which were then implanted into pseudopregnant mice. As the initial litter of mice likely consisted of chimeras, the F1 generation was crossed with each other to generate a non-chimeric F2 generation. The F2 generation was then sequenced, confirming the presence of homozygous *Enpp1^H362A^* mutations in several mice.

### cGAMP ELISA for basal cGAMP detection

Mice were euthanized and the spleens, kidneys, livers, and lungs were harvested. The spleens were diluted with 7.5 mL/g PBS and all other tissues were diluted with 2.5 mL/g PBS. The tissues were then homogenized and spun down at 2,000 x *g* for 15 min. A commercial cGAMP ELISA (Cayman Chemical) was used to determine the cGAMP concentration in each sample following the manufacturer’s specifications.

### Histology sectioning and staining

Organs were harvested at 20 weeks of age and fixed in 4% buffered Formaldehyde solution (pH 6.9) for 72 hours before transfer into 70% ethanol. Samples were submitted to Stanford Animal Histology Services for paraffin embedding, cutting and Alizarin Red staining. Imaging was done on a Zeiss AxioImager microscope in the Stanford Cell Sciences Imaging Facility.

### Plasma chemistry

Blood was collected through terminal cardiac puncture into heparin-coated microtainers (BD). The blood was then spun at 2,000 x g for 15 min, and the resulting plasma layer was collected. Plasma phosphate was measured using a malachite green phosphate assay kit (Sigma-Aldrich) according to the manufacturer’s instructions; each sample was diluted 1:250 in water. Plasma calcium was measured using a commercial colorimetric assay (Stanbio) according to the manufacturer’s instructions; each sample was diluted 1:4 in water. Plasma pyrophosphate was measured using a previously published method (Oheim et al., 2020). To convert pyrophosphate into ATP, 5 μL of each plasma sample was added to a reaction mixture consisting of 16 μM adenosine phosphosulfate, 80 mM MgSO_4_, 50 mM HEPEs, and 0.5 µL ATP sulfurylase (MCLab). The reaction mixture was then incubated at 37°C for 10 min, followed by 90°C for 10 min to inactivate the enzyme. In order to measure ATP, 25 μL of the reaction mixture was added to 25 μL of CellTiter-Glo (Promega). Luminescence was measured after 10 min using a 0.5 s integration time.

### Transfecting and harvesting cells with ENPP1 plasmids

Plasmids containing Flag-ENPP1 WT or ENPP1 mutations were transfected into 293T *ENPP1*^-/-^ cells with polyethylenimine (PEI) at ratio of 1 μg plasmid to 3 μg PEI per well of a 12 well plate. After 24 hours, cells were lysed for western blotting and activity assays. For western blotting, cells were lysed on the plate in 150 μL of Laemmli sample buffer. For activity assays, cells were washed off the plate in 1 mL of PBS, centrifuged at 1000 x *g* for 10 minutes, and lysed in 100 μL of lysis buffer (10 mM Tris pH 9, 150 mM NaCl, 10 μM ZnCl_2_, 1% NP-40). Lysates were stored at −20 °C.

### *In vivo* cGAMP metabolism

Mice were injected subcutaneously with 5 mg/kg cGAMP diluted in 100 μL PBS. After 30 min, the mice were anesthetized with isoflurane and 50 μL of blood was collected retro-orbitally and immediately supplemented with ~20 μM of ENPP1 inhibitor STF-1623 (Carozza et al., 2020b) to stop cGAMP degradation by ENPP1 post-draw. The blood was placed in heparin-coated microtainers (BD) and spun at 2,000 x g for 15 min, and the resulting plasma layer was collected. The plasma was processed for LC-MS/MS by mixing plasma (7 μL) with acetonitrile containing 2 μM of internal standard cyclic GMP-[^13^C_10_,^15^N_5_]AMP (20 μL) (Carozza et al., 2020a), centrifuging at 16,000 x *g* for 15 min, and then adding 23 μL of the mixture to 15 μL of water containing 0.1% formic acid. cGAMP was analyzed on a Q-Exactive FT mass spectrometer (Thermo) equipped with a Vanquish UHPLC. Samples were injected onto a Biobasic AX LC column (5 μm, 50 × 3 mm; Thermo Scientific). The mobile phase consisted of 100 mM ammonium carbonate (A) and 0.1% formic acid in acetonitrile (B). The initial condition was 90% B, maintained for 0.5 min. The mobile phase was ramped to 30% A from 0.5 min to 2.0 min, maintained at 30% A from 2.0 min to 3.5 min, ramped to 90% B from 3.5 min to 3.6 min and maintained at 90% B from 3.6 min to 5 min. The flow rate was set to 0.6 mL min^-1^. Quantification was performed with TraceFinder 4.1 software (Thermo Fisher).

### HSV-1 purification

The HSV-1 KOS strain was purchased from ATCC. The day prior to infection, Vero cells were plated in five T175 tissue culture flasks (Corning) at a density of 8 x 10^6^ cells/flask so they would be 80-100% confluent on the day of infection. Cells were infected with HSV-1 at MOI 0.01 in 5 mL/flask of serum-free DMEM for 1 hour with gentle rocking every 15 minutes. Media was collected 48 hours post infection when the cells displayed >90% CPE and centrifuged at 600 x *g* for 10 min to pellet debris. Clarified media was then centrifuged at 48,000 x *g* for 30 min to pellet virus. The pelleted virus was gently washed with PBS and resuspended in 2 mL of PBS, aliquoted, snap frozen, and stored at −80 °C until further use. Plaque assay was performed to determine titer (usually ~1 x 10^9^ pfu/mL).

### *In vivo* HSV-1 infection model

2.5 x 10^7^ PFU of HSV-1 was diluted in 100 μL PBS and injected intravenously into the tail vein of each mouse. After 12 h, the mice were euthanized in a CO_2_ chamber and blood and organs were harvested. The blood was collected through cardiac puncture into heparin-coated microtainers (BD). The blood was then spun at 2,000 x g for 15 min, and the resulting plasma layer was collected. The organs were placed into collection tubes and frozen at −80°C until further processing.

### Plaque assays

One day prior to infection, Vero cells were plated at 0.2 x 10^6^ cells/well in a 12-well plate or 0.1 x 10^6^ cells/well in a 24-well plate so they would be 80-100% confluent on the day of infection. For tittering of HSV-1 stocks, 10-fold dilutions of HSV-1 were prepared in serum-free DMEM. For tittering of HSV-1 from tissues, previously frozen tissues were homogenized with 2.0 mm disruption beads (RPI) in Sarstedt tubes (Fisher, 50-809-242) using a tissue homogenizer. Final concentration of tissue homogenates was 500 mg/mL in PBS or serum-free DMEM. Tissue homogenates were centrifuged at 500 x *g* for 5 minutes, and 2-fold and 20-fold dilutions were prepared in serum-free DMEM. To infect, Vero cells were washed one time with PBS, infected with 100 μL of sample for 1 hour with gentle rocking every 15 minutes, and then overlaid with complete DMEM containing 10 μg/mL human IgG (Sigma). 48 hours post infection, cells were fixed in 10% paraformaldehyde and stained with 0.4% crystal violet in 20% methanol.

### RT-qPCR

Total RNA was isolated from cells and tissues using TRIzol (Invitrogen) by following the manufacturer’s protocol. Tissue samples were homogenized in TRIzol prior to RNA isolation. To obtain cDNA, 20 uL reverse transcriptase (RT) reactions were set up containing 1 μg total RNA, 100 pmol random hexamer primers, 0.5 mM dNTPs, 20 U RNaseOUT, 1x Maxima RT buffer, and 200 U Maxima RT (Thermo Scientific). RT reactions were incubated for 10 min at 25 °C, 15 min at 50 °C, then 5 min at 85 °C. To measure transcript levels, 10 uL qPCR reactions were set up containing 0.7 uL cDNA, 100 nM qPCR primers (**Supplementary Table 1**), and 1x AccuPower GreenStar master mix (Bioneer) or 1x PowerTrack SYBR Green master mix (Thermo Scientific). Reactions were run on a ViiA 7 Real-Time PCR System (Applied Biosystems) using the following program: ramp up to 50°C (1.6°C/s) and incubate for 2 min, ramp up to 95°C (1.6°C/s) and incubate for 10 min, then 40 cycles of the following: ramp up to 95°C (1.6°C/s) and incubate for 15 s, then ramp down to 60°C (1.6°C/s) and incubate for 1 min. Transcript levels for each gene were normalized to *Actb* transcript levels.

### cGAMP ELISA for plasma cGAMP measurement

2.5 x 10^7^ PFU of HSV-1 was diluted in 100 μL PBS and injected intravenously into the tail vein of each mouse. After the indicated timepoints, the mice were euthanized in a CO_2_ chamber the blood was collected through cardiac puncture into heparin-coated microtainers (BD). The blood was then spun at 2,000 x g for 15 min, and the resulting plasma layer was collected. A commercial cGAMP ELISA (Cayman Chemical) was used to determine the cGAMP concentration in each sample following the manufacturer’s specifications. Each sample was diluted 1:2 in the provided buffer, and the standard curve was generated in buffer mixed with 50% mouse plasma from uninfected *Enpp1^H362A^* mice.

### Statistical analysis

All statistical tests were performed using GraphPad Prism software and are noted in the figure legends. Data are presented as the mean ± standard deviation unless otherwise stated.

## Data and Materials Availability

All data are available in the main text or the supplementary materials.

## Acknowledgements

We thank S. Wang for signal peptide analysis, F. Sunden for *Xac* NPP plasmids and expression advice, and K. Nguyen for LC-MS/MS analysis. We thank all Li Lab members for their constructive comments and discussion through the course of this study. We thank the Stanford Transgenic, Knockout, and Tumor Center for their assistance in generating transgenic mice for this project. J.A.C. was supported by NIH 5F31CA239510 and the Stanford Interdisciplinary Graduate Fellowship affiliated with ChEM-H. A.F.C. was supported by NIH 5T32GM736544 and 1F30CA250145. This work was supported by NIH DP2CA228044 (L.L).

## Author Contributions

J.A.C., A.F.C, and L.L. designed the study. J.A.C., A.F.C, Y.A., V.B, and G.S. performed experiments. J.A.C., A.F.C, and L.L. wrote the manuscript. All authors discussed the findings and commented on the manuscript.

