## Supplemental Information for "Probing pathophysiology of extracellular cGAMP with substrate-selective ENPP1"

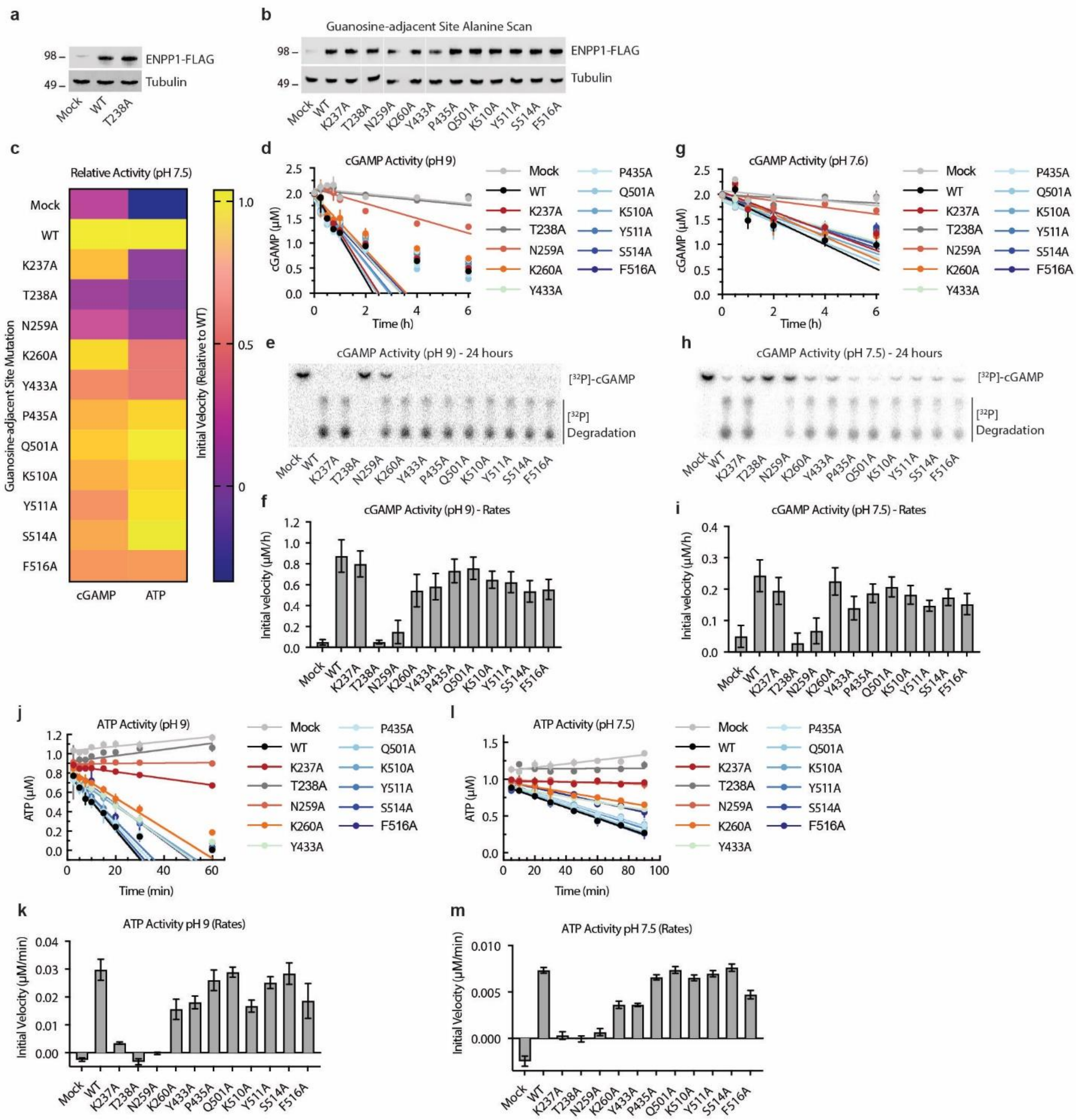


**Supplementary Fig. 1 Kinetic analysis and expression of ENPP1 guanosine-binding pocket mutations. a** Expression of $ENPP1^{WT}$ and $ENPP1^{T238A}$ mutation, representative of 3 independent experiments. **b** Expression of ENPP1 guanosine-binding pocket mutations, representative of 2 independent experiments. **c** Heat map showing the initial velocity relative to WT of guanosine-binding pocket mutations for the substrates cGAMP and ATP at pH 7.5. **d-i** cGAMP degradation kinetics of ENPP1 guanosine-binding pocket mutations at pH 9 (**d-f**) and pH 7.5 (**g-i**). Cell lines were transfected with the indicated ENPP1 mutations. Lines in (**d**) and (**g**) represent linear fits of kinetic data during the linear portion of the reaction. Time = 0 to 1 h for (**d**) and time = 0 to 6 h for (**g**). Data is representative of 2 independent experiments. These linear fits were plotted in bar graphs in (**f**) and (**i**) showing the best fit slope ± SD of the fit and were used to make heat maps shown in Fig. 1h and (**c**). TLCs in (**e**) and (**h**) depict reaction progress after 24 hours. **j-m** ATP degradation kinetics of ENPP1 guanosine-binding pocket mutations at pH 9 (**j-k**) and pH 7.5 (**l-m**). Cell lines were transfected with the indicated ENPP1 mutations. Lines in (**j**) and (**l**) represent linear fits of kinetic data during the linear portion of the reaction. Time = 0 to 20 min for (**j**) and time = 0 to 90 min for (**l**). $n$ = 4 independent reactions, mean ± SD shown with some error bars too small to visualize. These linear fits were plotted in bar graphs in (**k**) and (**m**) showing the best fit slope ± SD of the fit and were used to make heat maps shown in Fig. 1h and (**c**).

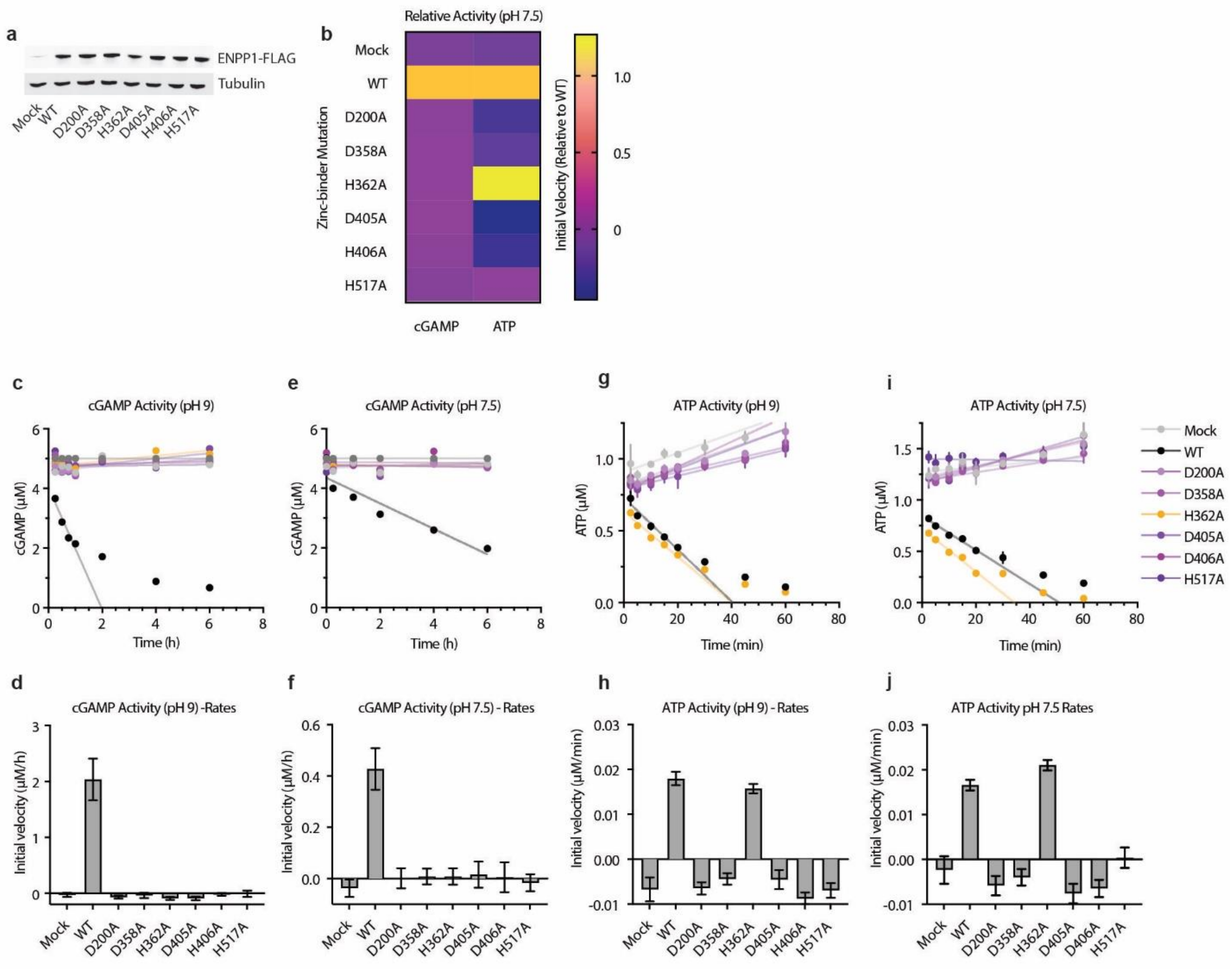


**Supplementary Fig. 2 Kinetic analysis and expression of ENPP1 zinc binder mutations.** **a** Expression of ENPP1 zinc-binding residue mutations, representative of 2 independent experiments. **b** Heat map showing the initial velocity relative to WT of zinc-binding residue mutations for the substrates cGAMP and ATP at pH 7.5. The mean initial velocity was calculated from a linear fit of the degradation reactions during early time points. *n* = 3 independent reactions. **c-f** cGAMP degradation kinetics of ENPP1 zinc-binding residue mutations at pH 9 (**c–d**) and pH 7.5 (**e–f**). Cell lines were transfected with the indicated ENPP1 mutations. Lines in (**c**) and (**e**) represent linear fits of kinetic data during the linear portion of the reaction. Time = 0 to 45 min for (**c**) and time = 0 to 6 h for (**e**). Data is representative of 2 independent experiments. These linear fits were plotted in bar graphs in (**d**) and (**f**) showing the best fit slope ± SD of the fit and were used to make heat maps shown in Fig. 2c and (**b**). **g-j** ATP degradation kinetics of ENPP1 zinc-binding residue mutations at pH 9 (**g-h**) and pH 7.5 (**i-j**). Cell lines were transfected with the indicated ENPP1 mutations. Lines in (**g**) and (**i**) represent linear fits of kinetic data during the linear portion of the reaction (time = 0 to 20 min). *n* = 3 independent reactions, mean ± SD with some error bars too small to visualize. These linear fits were plotted in bar graphs in (**h**) and (**j**) showing the best fit slope ± SD of the fit and were used to make heat maps shown in Fig. 2c and (**b**).

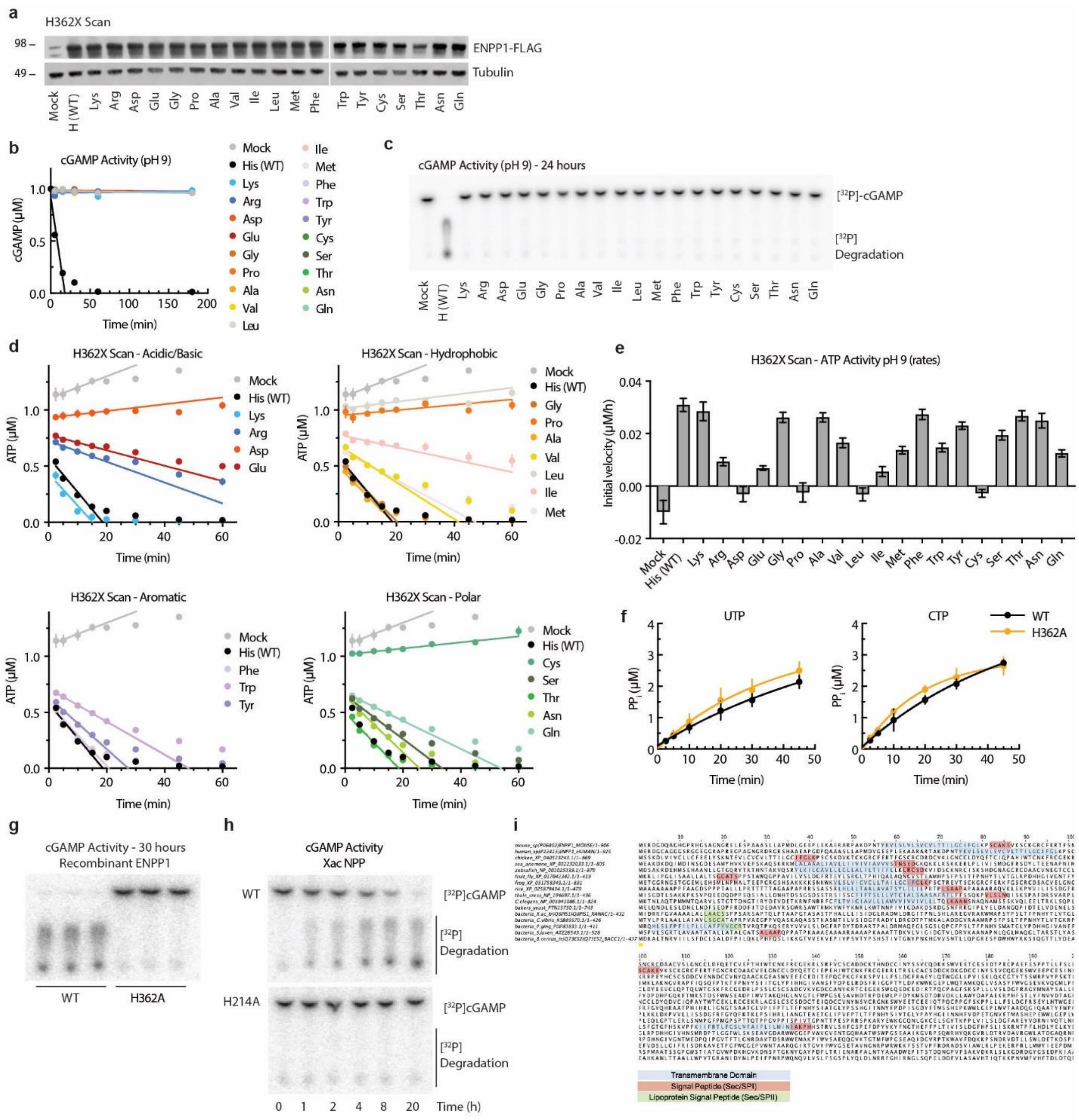


**Supplementary Fig. 3 Kinetic analysis and expression of ENPP1 H362X mutations. a** Expression of ENPP1 H362X mutations (where X represents the indicated amino acid), representative of 2 independent experiments. **b** cGAMP degradation kinetics of ENPP1 H362X mutations at pH 9. Cell lines were transfected with the indicated ENPP1 mutations. Lines represent linear fits of kinetic data during the linear portion of the reaction (time = 0 to 30 minutes). **c** TLC of cGAMP degradation of ENPP1 H362X mutations after 24 hours at pH 9. **d** ATP degradation kinetics of ENPP1 H362X mutations at pH 9, organized by amino acid class. Cell lines were transfected with the indicated ENPP1 mutations. Lines represent linear fits of kinetic data during the linear portion of the reaction (time = 0 to 15 min). *n* = 3 independent reactions, mean ± SD with some error bars too small to visualize. **e** ATP activity rates for ENPP1 H362X mutations plotted from linear fits shown in (**d**). *n* = 3 independent reactions, best fit slope ± SD of the fit from (**d**). **f** Kinetic analysis of UTP and CTP monitoring pyrophosphate production using recombinant purified ENPP1. *n* = 2 independent reactions, mean ± SD. **g** TLC of cGAMP degradation comparing recombinant ENPP1$^{WT}$ to ENPP1$^{H362A}$. *n* = 3 independent reactions. **h** TLC of cGAMP degradation comparing *Xac* NPP$^{WT}$ to *Xac* NPP$^{H214A}$ (both at 1 μM enzyme concentration). Representative of 3 independent reactions. **i** Signal peptides (red boxes) or lipoprotein signal peptides (blue boxes) for selected species predicted by SignalP5.0. No signal peptide was found for *B. cereus*. Baker's yeast, sea anemone, and fruit fly are shown on a separate alignment since their signal peptide regions aligned poorly with the rest of the eukaryotic sequences.

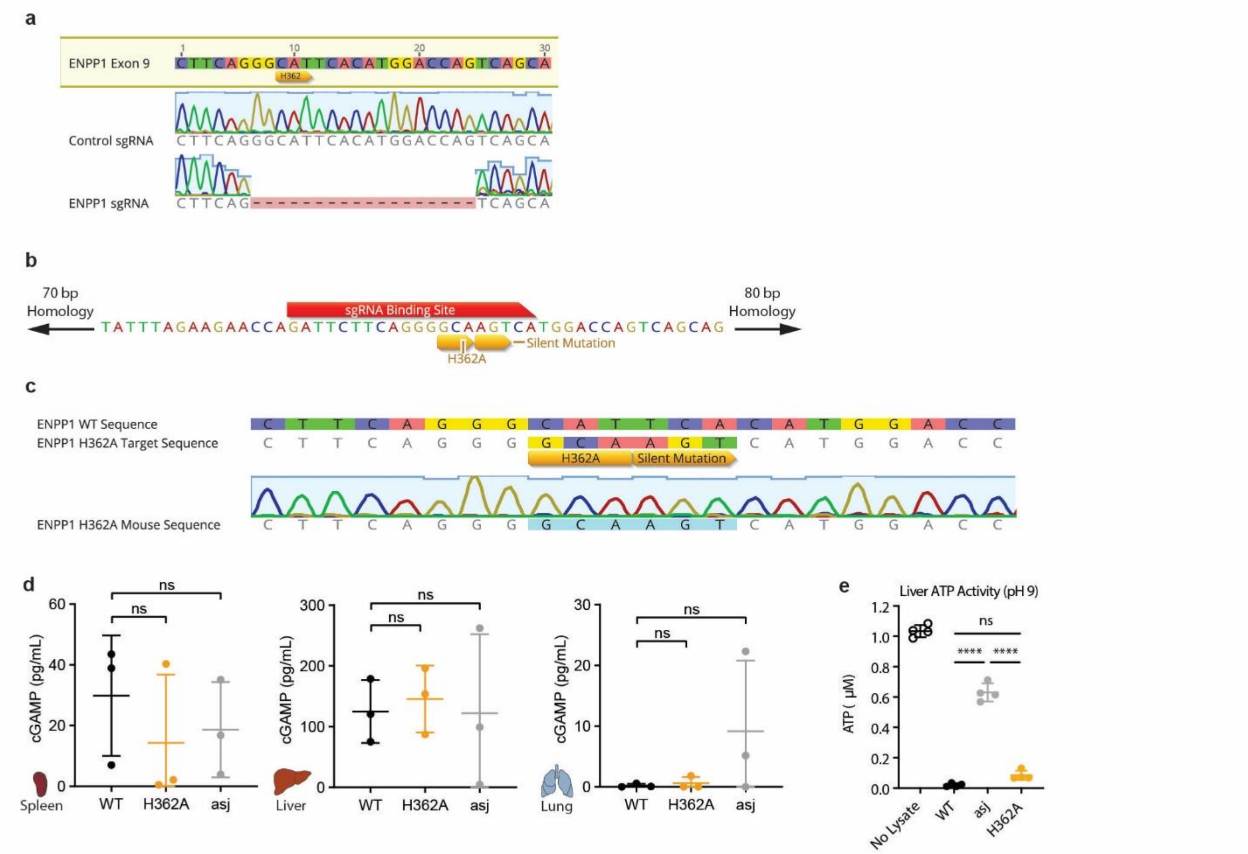


**Supplementary Fig. 4 *Enpp1$^{H362A}$* mice can degrade ATP but not cGAMP. a** A single-guide RNA (sgRNA) was designed to target the region near H362A in exon 9 of Enpp1. The sgRNA and Cas9 were introduced into HEK 239T cells through lentiviral transduction. The cells were then sequenced to determine editing efficiency at the intended cleavage site. There was no evidence of editing in cells transduced with a control sgRNA, while there was a large deletion in cells transduced with the *Enpp1* sgRNA. **b** Diagram indicating the sequence of the donor ssDNA used to generate the H362A point mutation through homologous recombination. Silent mutations were introduced downstream of H362 to prevent Cas9 from recognizing the cleavage site following successful recombination. **c** Genomic DNA sequencing from one of the *Enpp1$^{H362A}$* mice. The sequencing indicated that this mouse harbored a homozygous H362A point mutation in *Enpp1*, as well as the point mutations indicated in (**b**). **d** A cGAMP ELISA was performed to measure basal cGAMP in the spleen, kidney, liver, and lung. There was no detectable cGAMP in any of the kidneys, so they were omitted from analysis. n = 3 mice per genotype. *p* values were calculated by unpaired *t* test; *$p < 0.05$. **e** *Ex vivo* liver lysate (1 mg/mL) ATP degradation at pH 9 assessed by luciferase assay after 20 minutes in *Enpp1$^{H362A}$*, *Enpp1$^{asj}$*, and *Enpp1$^{H362A}$* mice. *n* = 4 *Enpp1$^{H362A}$*, 4 *Enpp1$^{asj}$*, and 4 *Enpp1$^{H362A}$* mice.

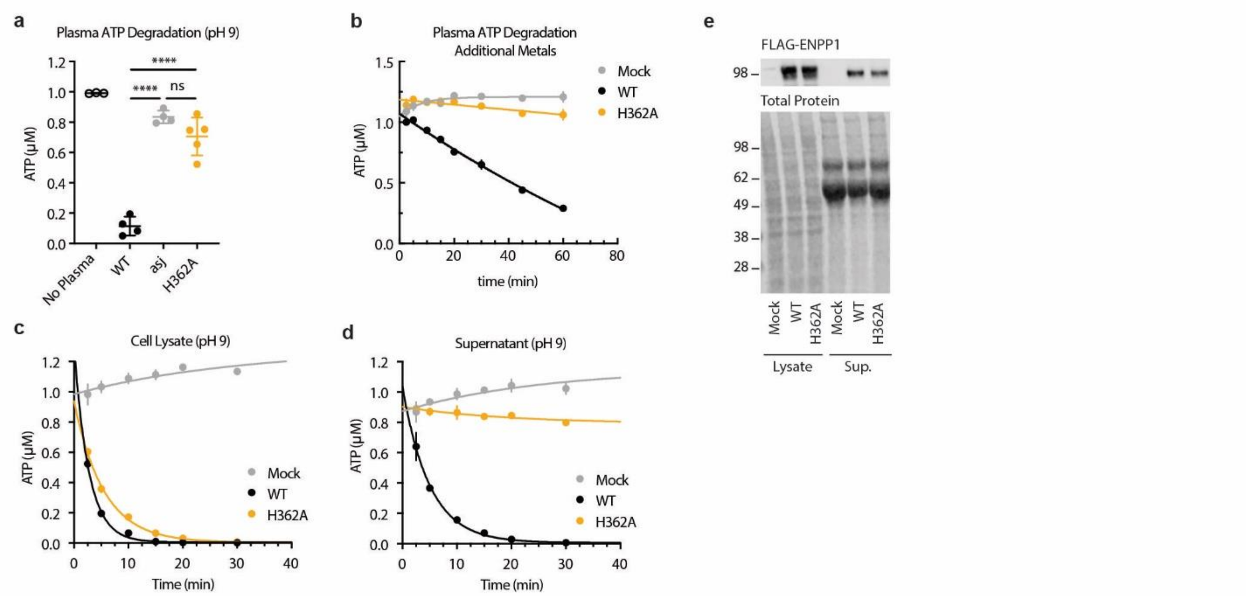


**Supplementary Fig. 5 *Enpp1$^{H362A}$* mice do not exhibit systemic calcification despite low plasma pyrophosphate. a** *Ex vivo* plasma ATP degradation at pH 9 assessed by luciferase assay after 45 minutes in *Enpp1$^{WT}$*, *Enpp1$^{asj}$*, and *Enpp1$^{H362A}$* mice. $n$ = 4 *Enpp1$^{WT}$*, 4 *Enpp1$^{asj}$*, and 5 *Enpp1$^{H362A}$* mice. Data are shown as the mean ± SD. $p$ values were calculated by unpaired $t$ test with Welch's correction. ****$p < 0.0001$. **b** *Ex vivo* plasma ATP degradation kinetics at pH 9 with additional metals (100 µM $ZnCl_2$ and 10 mM $CaCl_2$) assessed by luciferase assay after 45 minutes in *Enpp1$^{WT}$*, *Enpp1$^{asj}$*, and *Enpp1$^{H362A}$* mouse plasma. **c-d** *In vitro* ATP degradation comparing overexpressed ENPP1$^{WT}$ and ENPP1$^{H362A}$ as cell-surface protein from cell lysate (**c**) and as secreted protein from cell supernatant (**d**) at pH 9. **e** Western blot of FLAG-ENPP1 as cell-surface protein from cell lysate and as secreted protein from cell supernatant. Supernatant was concentrated 16-fold prior to blotting for clear visualization.

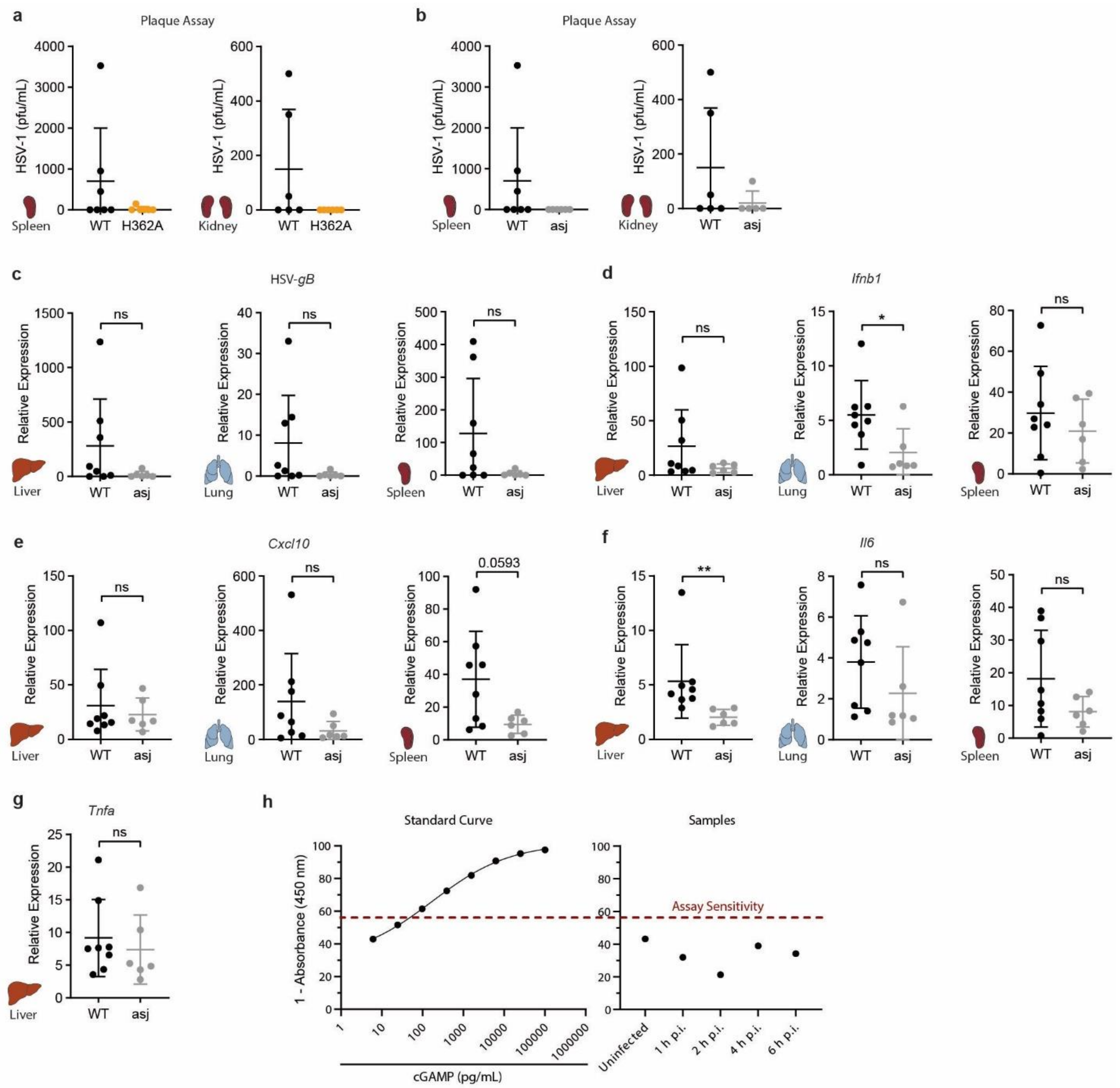


**Supplementary Fig. 6 *Enpp1*$^{H362A}$ mice are resistant to HSV-1 infection. a-g** Mice were inoculated with 2.5 x $10^7$ PFU/mouse HSV-1 through intravenous injection. The mice were euthanized after 12 h, and organs were isolated for plaque assays or total RNA isolation. n = 8 infected *Enpp1*$^{WT}$ and *ENPP1*$^{H362A}$ mice (as previously described in Fig. 5) and n = 4 *Enpp1*$^{asj}$ mice. One *Enpp1*$^{asj}$ mouse was removed as an outlier based on the ROUT method (Q = 1%). **a-b** Plaque assays comparing spleen and kidney lysates from *Enpp1*$^{WT}$ and *Enpp1*$^{H362A}$ mice (**a**) or *Enpp1*$^{WT}$ and *Enpp1*$^{asj}$ mice (**b**). **c-g** RT-qPCR was performed to determine the expression levels of HSV-*gB* (**c**), *Ifnb1* (**d**), *Cxcl10* (**e**), *Il6* (**f**), and *Tnfa* (**g**). Transcript levels were normalized to the average of 2 uninfected mice per genotype. Data are shown as the mean ± SD. p values were calculated using the Mann-Whitney test. **p* < 0.05, ***p* < 0.01; p value is shown if between 0.05 and 0.1. **h** *Enpp1*$^{H362A}$ mice were injected with 2.5 x $10^7$ PFU/mouse HSV-1 and euthanized at the indicated time points. Plasma was collected from each mouse, and cGAMP concentration was determined by a cGAMP ELISA. A cGAMP standard curve was created in 50% mouse plasma (left). None of the infected plasma samples (right) gave readings above the published limit of detection (85 pg/mL), suggesting the absence of any cGAMP.

**Supplemental Table 1. Oligonucleotides used in this study**

| **Primers for Cloning:** | |
|---|---|
| **Name** | **Sequence (5’->3’)** |
| mENPP1_D200A_fwd | CCCCCTACTCTCTTGTTTTCTTTGGCTGGATTCAGAGCTG |
| mENPP1_D200A_rev | CAGCTCTGAATCCAGCCAAAGAAAACAAGAGAGTAGGGGG |
| mENPP1_K237A_fwd | CCTATGTACCCTACCGCAACGTTTCCCAATCATTACAGC |
| mENPP1_K237A_rev | GCTGTAATGATTGGGAAACGTTGCGGTAGGGTACATAGG |
| mENPP1_T238A_fwd | GCCTATGTACCCTACCAAGgcgTTTCCCAATCATTACAGC |
| mENPP1_T238A_rev | GCTGTAATGATTGGGAAAcgcCTTGGTAGGGTACATAGGC |
| mENPP1_N259A_fwd | CCCATGGCATAATTGATGCAAAGATGTATGATCCC |
| mENPP1_N259A_rev | GGGATCATACATCTTTGCATCAATTATGCCATGGG |
| mENPP1_K260A_fwd | CCCATGGCATAATTGATAACGCAATGTATGATCCCAAAATGAATGC |
| mENPP1_K260A_rev | GCATTCATTTTGGGATCATACATTGCGTTATCAATTATGCCATGGG |
| mENPP1_D358A_fwd | GTATTTAGAAGAACCAGcTTCTTCAGGGCATTCACATGG |
| mENPP1_D358A_rev | CCATGTGAATGCCCTGAAGAAgCTGGTTCTTCTAAATAC |
| mENPP1_H362A_fwd | CCAGATTCTTCAGGGGCATCACATGGACCAGTCAGC |
| mENPP1_H362A_rev | GCTGACTGGTCCATGTGATGCCCCTGAAGAATCTGG |
| mENPP1_D405A_fwd | CCTCATCCTCATTTCAGcTCACGGCATGGAACAAGG |
| mENPP1_D405A_rev | CCTTGTTCCATGCCGTGAgCTGAAATGAGGATGAGG |
| mENPP1_H406A_fwd | CATCCTCATTTCAGATgcCGGCATGGAACAAGGCAGC |
| mENPP1_H406A_rev | GCTGCCTTGTTCCATGCCGgcATCTGAAATGAGGATG |
| mENPP1_Y433A_fwd | GGATGTGAACAATGTGAAAGTTGTGGCAGGACCTGCTGCTCGG |
| mENPP1_Y433A_rev | CCGAGCAGCAGGTCCTGCCACAACTTTCACATTGTTCACATCC |
| mENPP1_P435A_fwd | GTGAAAGTTGTGTATGGAGCAGCTGCTCGGTTGAGACCC |
| mENPP1_P435A_rev | GGGTCTCAACCGAGCAGCTGCTCCATACACAACTTTCAC |
| mENPP1_Q501A_fwd | CCTGGACCCTCAGTGGGCACTTGCGTTGAATCCATCAGAGAGG |
| mENPP1_Q501A_rev | CCTCTCTGATGGATTCAACGCAAGTGCCCACTGAGGGTCCAGG |
| mENPP1_K510A_fwd | CCATCAGAGAGGGCATATTGTGGAAGTGG |
| mENPP1_K510A_rev | CCACTTCCACAATATGCCCTCTCTGATGG |
| mENPP1_Y511A_fwd | CCATCAGAGAGGAAAGCATGTGGAAGTGGATTTCATGG |
| mENPP1_Y511A_rev | CCATGAAATCCACTTCCACATGCTTTCCTCTCTGATGG |
| mENPP1_S514A_fwd | GGAAATATTGTGGAGCAGGATTTCATGGCTCTGAC |
| mENPP1_S514A_rev | GTCAGAGCCATGAAATCCTGCTCCACAATATTTCC |
| mENPP1_F516A_fwd | GGAAATATTGTGGAAGTGGAGCACATGGCTCTGACAAC |
| mENPP1_F516A_rev | GTTGTCAGAGCCATGTGCTCCACTTCCACAATATTTCC |
| mENPP1_H517A_fwd | GTGGAAGTGGATTTgcTGGCTCTGACAACTTG |
| mENPP1_H517A_rev | CAAGTTGTCAGAGCCAgcAAATCCACTTCCAC |
| mENPP1_H362R_fwd | CCAGATTCTTCAGGGcgtTCACATGGACCAGTCAGC |
| mENPP1_H362R_rev | GCTGACTGGTCCATGTGAacgCCCTGAAGAATCTGG |
| mENPP1_H362E_fwd | CCAGATTCTTCAGGGgagTCACATGGACCAGTCAGC |
| mENPP1_H362E_rev | GCTGACTGGTCCATGTGActcCCCTGAAGAATCTGG |
| mENPP1_H362S_fwd | CCAGATTCTTCAGGGtctTCACATGGACCAGTCAGC |
| mENPP1_H362S_rev | GCTGACTGGTCCATGTGAagaCCCTGAAGAATCTGG |
| mENPP1_H362T_fwd | CCAGATTCTTCAGGGactTCACATGGACCAGTCAGC |
| mENPP1_H362T_rev | GCTGACTGGTCCATGTGAagtCCCTGAAGAATCTGG |

| | |
|---|---|
| mENPP1_H362N_fwd | CCAGATTCTTCAGGGaatTCACATGGACCAGTCAGC |
| mENPP1_H362N_rev | GCTGACTGGTCCATGTGAattCCCTGAAGAATCTGG |
| mENPP1_H362C_fwd | CCAGATTCTTCAGGGtgtTCACATGGACCAGTCAGC |
| mENPP1_H362C_rev | GCTGACTGGTCCATGTGAacaCCCTGAAGAATCTGG |
| mENPP1_H362G_fwd | CCAGATTCTTCAGGGggtTCACATGGACCAGTCAGC |
| mENPP1_H362G_rev | GCTGACTGGTCCATGTGAaccCCCTGAAGAATCTGG |
| mENPP1_H362P_fwd | CCAGATTCTTCAGGGcctTCACATGGACCAGTCAGC |
| mENPP1_H362P_rev | GCTGACTGGTCCATGTGAaggCCCTGAAGAATCTGG |
| mENPP1_H362I_fwd | CCAGATTCTTCAGGGattTCACATGGACCAGTCAGC |
| mENPP1_H362I_rev | GCTGACTGGTCCATGTGAaatCCCTGAAGAATCTGG |
| mENPP1_H362M_fwd | CCAGATTCTTCAGGGatgTCACATGGACCAGTCAGC |
| mENPP1_H362M_rev | GCTGACTGGTCCATGTGAcatCCCTGAAGAATCTGG |
| mENPP1_H362W_fwd | CCAGATTCTTCAGGGtggTCACATGGACCAGTCAGC |
| mENPP1_H362W_rev | GCTGACTGGTCCATGTGAccaCCCTGAAGAATCTGG |
| mENPP1_H362Y_fwd | CCAGATTCTTCAGGGtatTCACATGGACCAGTCAGC |
| mENPP1_H362Y_rev | GCTGACTGGTCCATGTGAataCCCTGAAGAATCTGG |
| mENPP1_H362V_fwd | CCAGATTCTTCAGGGgttTCACATGGACCAGTCAGC |
| mENPP1_H362V_rev | GCTGACTGGTCCATGTGAaacCCCTGAAGAATCTGG |
| mENPP1_seq_rev | CCATTATTGGGAGCTGGGATCAAACC |
| mENPP1_seq_fwd | CTACAGTTCTGTGTGCCAAG |
| XacNPP_H214A_fwd | CATGTGGACGAAGCCGGCgcCGACCACGGCCCGGAATCGC |
| XacNPP_H214A_rev | GCGATTCCGGGCCGTGGTCGgcGCCGGCTTCGTCCACATG |
| **Primers for qPCR:** | |
| **Name** | **Sequence (5’->3’)** |
| Cxcl10 Fwd | AAGTGCTGCCGTCATTTTCT |
| Cxcl10 Rev | GTGGCAATGATCTCAACACG |
| Irf7 Fwd | GAAGACCCTGATCCTGGTGA |
| Irf7 Rev | CCAGGTCCATGAGGAAGTGT |
| HSV-gB Fwd | ATTCTCCTCCGACGCCATATCCACCACCTT |
| HSV-gB Rev | AGAAAGCCCCCATTGGCCAGGTAGT |
| Actb Fwd | AGCCATGTACGTAGCCATCC |
| Actb Rev | CTCTCAGCTGTGGTGGTGAA |
| **Oligonucleotides for generating the ENPP1$^{H362A}$ mouse:** | |
| **Name** | **Sequence (5’->3’)** |
| Enpp1 Exon 9 sgRNA | GATTCTTCAGGGCATTCACA |
| Enpp1 Exon 9 Sequencing Fwd | GATGATTTATAGCCAGAGCAACTAGTG |
| Enpp1 Exon 9 Sequencing Rev | GTTCTCTCTGGCTACATAGAATTTC |
| Enpp1 H362A Donor Sequence for Homologous Recombination | TGTTTTTCAATGTGTTCGTAAAATGTTACATTTTGATACTGTT TGATTTAGACCACACTTTTACACTCTGTATTTAGAAGAACCA GATTCTTCAGGGGCAAGTCATGGACCAGTCAGCAGCGAGG TAAGTTCACCGCTACCTATAATCACTTCGTTAAATTTAGTATT |

| | |
|---|---|
| | CCTGAAGTGGACTTCAAGAACCTTCTGTAAGG |
